# *Spiroplasma* endosymbiont reduction of host lipid synthesis and Stomoxyn-like peptide contribute to trypanosome resistance in the tsetse fly *Glossina fuscipes*

**DOI:** 10.1101/2024.10.24.620045

**Authors:** Erick Awuoche, Gretchen Smallenberger, Daniel Bruzzese, Alessandra Orfano, Brian L. Weiss, Serap Aksoy

**Affiliations:** Department of Epidemiology of Microbial Diseases, Yale School of Public Health, New Haven, CT, USA

**Author notes:** Northeast Scientific Inc. 2142 Thomaston Ave, Waterbury, CT 06704.

**Keywords:** *Spiroplasma*, Stomoxyn, vector competence, trypanosome, tsetse, *Glossina fuscipes*, lipids

## Abstract

Tsetse flies (*Glossina* spp.) vector African trypanosomes that cause devastating diseases in humans and domestic animals. Within the *Glossina* genus, species in the Palpalis subgroup exhibit greater resistance to trypanosome infections compared to those in the *Morsitans* subgroup. Varying microbiota composition and species-specific genetic traits can significantly influence the efficiency of parasite transmission. Notably, infections with the endosymbiotic bacterium *Spiroplasma* have been documented in several Palpalis subgroup species, including *Glossina fuscipes fuscipes* (*Gff*). While *Spiroplasma* infections in *Gff* are known to hinder trypanosome transmission, the underlying mechanisms remain unknown. To investigate *Spiroplasma-*mediated factors affecting *Gff* vector competence, we conducted high-throughput RNA sequencing of the midgut tissue along with functional assays. Our findings reveal elevated oxidative stress in the midgut environment in the presence of *Spiroplasma*, evidenced by increased expression of *nitric oxide synthase,* which catalyzes the production of trypanocidal nitric oxide. Additionally, we observed impaired lipid biosynthesis leading to a reduction of this important class of nutrients essential for parasite and host physiologies. In contrast, trypanosome infections in *Gff’s* midgut significantly upregulated various immunity-related genes, including a small peptide, *Stomoxyn-like*, homologous to Stomoxyns first discovered in the stable fly *Stomoxys calcitrans*. We observed that the *Stomoxyn-like* locus is exclusive to the genomes of *Palpalis* subgroup tsetse species. *Gff*Stomoxyn is constitutively expressed in the cardia (proventriculus) and synthetic *Gff*Stomoxyn exhibits potent activity against *Escherichia coli* and bloodstream form of *Trypanosoma brucei* parasites, while showing no effect against insect stage procyclic forms or tsetse’s commensal endosymbiont *Sodalis in vitro*. Reducing *Gff*Stomoxyn levels significantly increased trypanosome infection prevalence, indicating its potential trypanocidal role *in vivo*. Collectively, our results suggest that the enhanced resistance to trypanosomes observed in *Spiroplasma*-infected *Gff* may be due to the reduced lipid availability necessary for parasite metabolic maintenance. Furthermore, *Gff*Stomoxyn could play a crucial role in the initial immune response(s) against mammalian parasites early in the infection process in the midgut and prevent gut colonization. We discuss the molecular characteristics of *Gff*Stomoxyn, its spatial and temporal expression regulation and its microbicidal activity against *Trypanosome* parasites. Our findings reinforce the nutritional influences of microbiota on host physiology and host-pathogen dynamics.

**Author Summary:** The tsetse fly, *Glossina fuscipes fuscipes* (*Gff*) is of high public health relevance. Gff exhibits strong innate resistance to trypanosomes, especially when infected with the endosymbiotic bacterium *Spiroplasma*. This study investigated how the bacterium *Spiroplasma* inside *Gff* enables them to be resistant to trypanosome infection. Our results indicate alterations in host lipid metabolism with reduction in levels of triglycerides, suggesting a potential metabolic barrier that limits the viability to parasite. In addition, we discovered a small peptide, stomoxyn, exclusively in *Gff* and related *Palpalis* tsetse species. We have shown that *Gff* synthetic Stomoxyn has antibacterial and antitrypanosomal properties and lowering Stomoxyn levels in *Gff* correlates with increased parasite prevalence. We suggest that strategies to increase *Spiroplasma* prevalence or enhance stomoxyn expression through paratransgenic approaches could be promising avenues for reducing trypanosomiasis transmission.

## Introduction

Tsetse flies (*Glossina* spp.) transmit African trypanosome parasites that cause sleeping sickness (Human African Trypanosomiasis, HAT) in humans and Nagana (Animal African Trypanosomiasis, AAT) in livestock [1]. Approximately 60 million people live in tsetse fly-infested areas in sub-Sahara and hence are at risk of contracting HAT, while AAT is rampant and results in significant loss of agricultural productivity among the farming communities in impoverished areas of the continent [2, 3]. No vaccines exist to prevent mammalian infections due to a process of antigenic variation by which the parasites sequentially express antigenically distinct surface coat proteins to evade vertebrate host immune responses [4]. Reduction of tsetse fly populations can be effective at curbing the disease, but both the challenges and cost of implementing vector control activities and re-infestation risk once the programs are abandoned reduce their efficacy [5]. Blocking or reducing the ability of parasite transmission through the fly has been entertained as an additional method to boost disease control efforts [6–8]. For successful development of such alternative biological methods, better knowledge is required on parasite-vector dynamics, parasite transmission biology and antiparasitic molecules that could interfere with parasite transmission through tsetse fly vector.

Tsetse flies exhibit innate resistance to infection with trypanosomes, with low infection prevalences reported in wild and experimentally colonized fly populations [9–12]. Various factors influence parasite transmission efficiency under experimental conditions, including fly age and nutritional status at time of exposure to the parasite, species/strain of the trypanosome studied and resident microbiota in the fly midgut. Among the four subgroups of *Glossina* - Fusca, Palpalis, Morsitans and Machadomia-species within the Morsitans subgroup, including the well studied *Glossina morsitans morsitans* (*Gmm*), typically exhibit greater susceptibility to trypanosome infections compared to those in the Palpalis subgroup (e.g., *Glossina fuscipes fuscipes (Gff*)*, Glossina tachinoides* (*Gt*), *Glossina palpalis palpalis* (*Gpp*) and *Glossina palpalis gambiensis (Gpg)*) [11, 13–15]. The Palpalis subgroup, also known as riverine group, is widely distributed in West and Central Africa, covering an estimated area of 6415 square kilometers and living in close association with human habitats [16, 17]. Fly species in this subgroup are highly relevant to public health as they serve as the primary vectors for trypanosomes responsible for chronic sleeping sickness in the regions they inhabit [18, 19].

In addition to ecological differences and host preferences, variations in vector competence between Morsitans and Palpalis subgroup flies may arise from species-specific genetic content, as evidenced by comparisons of whole-genome sequencing (WGS) data across different *Glossina* subgroups [20]. A comparative analysis of orthology groups (OGs) among the four subgroups revealed the presence of 2223 OGs specific to the Palpalis subgroup, with 4948 genes shared between *Gff* and *Gpp* [20]. Notably the Palpalis subgroup exhibited gene expansions, including those encoding helicases involved in the production of small RNAs that mediate post-transcriptional gene expression and defensive responses against viruses and transposable elements, alongside gene duplications such as the trypanocidal peptide Cecropins in *Gff* [20, 21]. Laboratory studies in *Gmm* have shown that when flies acquire the bloodstream form (BSF) trypanosomes as newly eclosed adults (teneral) in their first bloodmeal, they exhibit higher susceptibility to parasite infections. However, older adults display greater midgut resistance and can eliminate the BSF trypanosomes early in the infection process in the midgut before they can colonize this organ. The trypanocidal factors described in *Gmm* include the gut peritrophic matrix [22–24], antimicrobial peptides [25–28], reactive oxygen species (ROS) [29, 30] tsetse EP proteins [31], trypanolysin [32–34], peptidoglycan recognition protein (PGRP)-LB [35], lectins [36–38] and other proteolytic enzymes [39–41]. Beyond physical barriers and innate immune factors, gut endosymbionts have also been implicated to influence parasite transmission success. Tsetse flies harbor a species-specific combination of four well-characterized microbes, including *Wigglesworthia*, *Sodalis*, *Wolbachia* and *Spiroplasma.* Each of these endosymbionts display a different evolutionary history with their vector species and exert varying influences on fly physiology. Several investigations have described a positive correlation between trypanosome infection prevalence and the presence of the commensal symbiont *Sodalis*, although the mechanism remains unconfirmed [10, 41]. The mutualist *Wigglesworthia* has been shown to induce the expression of a host amidase with trypanolytic activity (PGRP-LB) that reduces parasite colonization success as an early response during the infection process in the midgut [35]. Infections with *Spiroplasma glossinidia* (*Spiroplasma*) were reported uniquely from the species within the Palpalis subgroup, including *Gff* [42]*, Gt,* and *Gpp* [43, 44]. In Uganda, *Spiroplasma* infections have been found to be restricted in prevalence to distinct *Gff* populations in the Northwest region of the country and infections persist stably over time and space with seasonality being one important factor in infection prevalences [45]. Our studies with a *Gff* laboratory line in which approximately 50% of adults are infected with S*piroplasma* showed a negative correlation between the presence of this bacterium and trypanosome infection success [45]. A similar observation on trypanosome infection success reduction was reported in natural *Gt* populations in West Africa [44].

The objective of this study was to acquire a better understanding of the mechanism(s) underpinning the enhanced parasite refractoriness noted in *Gff*, including intrinsic fly specific and *Spiroplasma* regulated factors. For our analyses, we used the *Gff* lab line with a heterogenous *Spiroplasma* infection prevalence, and performed high-throughput RNA sequencing of midgut tissue collected from *Spiroplasma* infected (*Spi*^+^), *Spiroplasma* uninfected (*Ctrl*) and only trypanosome (no *Spiroplasma*) infected (*Tpi*^+^) individuals. We describe the host immune and metabolic responses elicited in the presence of *Spiroplasma* and trypanosomes and incriminate them as factors that limit parasite colonization success. We also discovered a small peptide uniquely present in the genomes of the tsetse species within the Palpalis subgroup. This peptide, designated *Gff*Stomoxyn is related to the Stomoxyn first described from the insect *Stomoxys calcitrans*. We investigated the spatial and temporal regulation of *GffStomoxyn* expression, its genomic context in tsetse species belonging to the different *Glossina* subgroups, and the microbicidal activity of *Gff*Stomoxyn against bacteria and BSF and insect stage procyclic (PCF) trypanosomes *in vitro.* We also report on the role of *Gff*Stomoxyn in trypanosome colonization success *in vivo*, via functional studies established through the use of dsRNA mediated RNAi. We discuss how *Spiroplasma* infection and Stomoxyn peptide may function to restrict parasite transmission processes in *Gff*.

## Results

We generated over one billion high quality reads obtained across 14 samples comprising five, five and four biological replicates that represent *Spiroplasma* infected (denoted as *Spi*^+^), *Trypanosome* infected (denoted as *Tpi*^+^), and uninfected (denoted as *Ctrl* with no *Spiroplasma* or trypanosome infection) groups, respectively (Fig. S1A). We obtained a Pearson correlation coefficient > 0.7 between the replicates in each experimental condition indicating that the libraries were of high related. Based on the predicted *Gff* transcriptome (*Gff* 2018 reference genome version 63), 84% of known genes were detected with ≥ 10 reads mapping in at least 50% of the biological replicates for each experimental condition (Fig. S1A; Table S2 Sheet 1 and 2). These mapping statistics suggested that the majority of the *Gff* transcripts were captured, giving us confidence for a robust downstream analysis.

### Tsetse midgut response to infection with symbiotic *Spiroplasma* or parasitic trypanosomes

To assess the impact of *Spiroplasma* or trypanosome infection outcomes on midgut functions, we performed transcriptional comparisons between the *Spi^+^* and *Tpi*^+^ groups relative to the uninfected *Ctrl* group. Infection with *Spiroplasma* resulted in a total of 69 differentially expressed (DE) genes, of which 49 were induced and 20 were suppressed (Fig. 1A; Table S2 Sheet 3). In contrast, infection with *Trypanosoma* resulted in a total of 989 DE genes, of which 518 were induced and 471 suppressed (Fig. 1A; Table S2 Sheet 4). Clustering based on the Euclidean distances between *Ctrl* and *Tpi^+^* samples resulted in a separate cluster group while no cluster groups were observed between *Ctrl* and *Spi^+^* samples (Fig. S1B). These results indicate that infections with trypanosomes induced a more robust response in the midgut in comparison to that elicited in the presence of *Spiroplasma* infections.

**Fig. 1.**
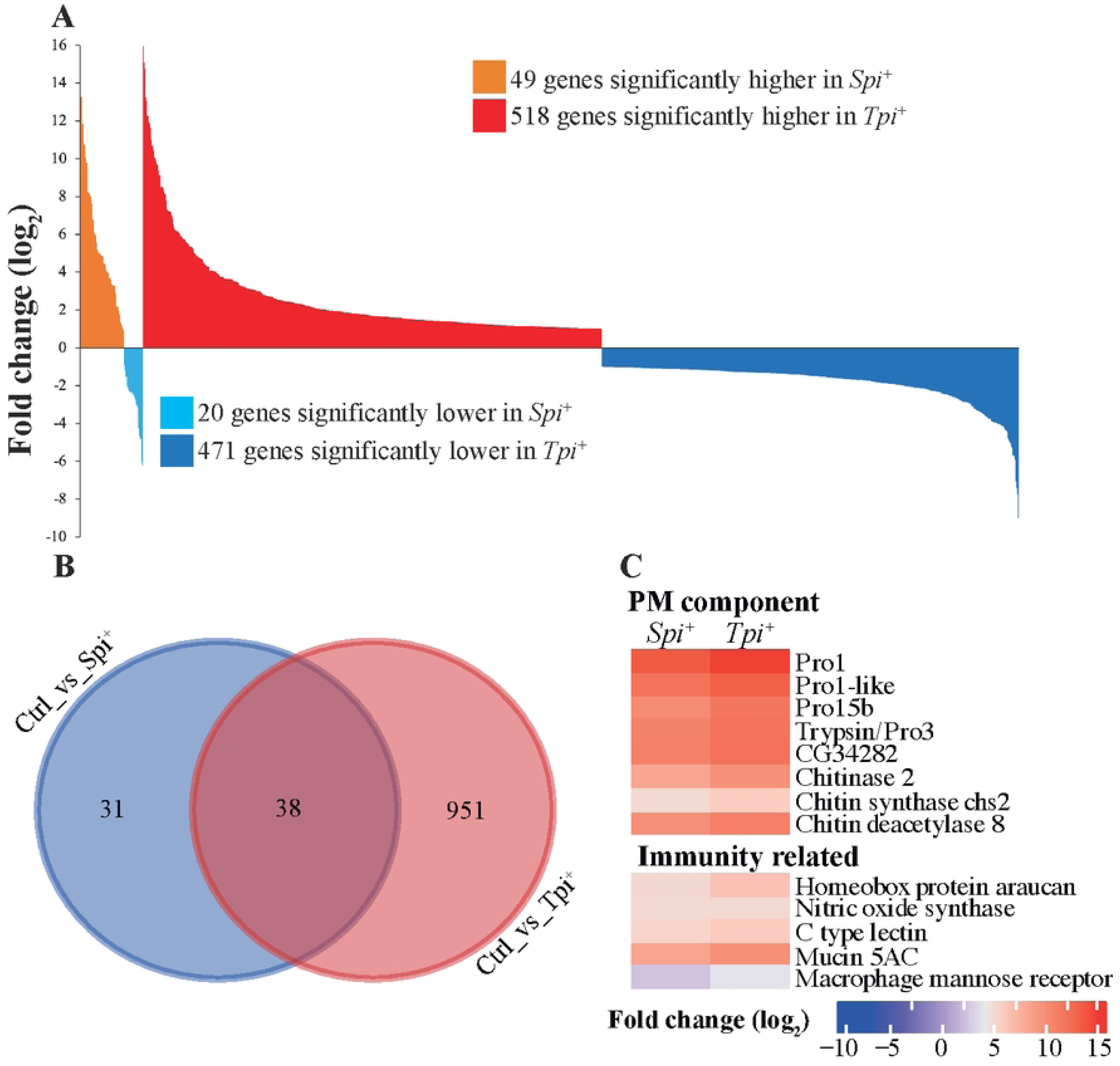
Tsetse Midgut Tissue Responses to *Spiroplasma* and Trypanosome Infection. **A**. Differentially expressed (DE) genes identified in the *Spi*^+^ and *Tpi*^+^ groups relative to uninfected *Ctrl* are presented as follows: The orange and red areas represent up-regulated genes in the *Spi*^+^ and *Tpi*^+^ state respectively, while light blue and dark blue indicate down-regulated genes in *Spi*^+^ and *Tpi*+ respectively, at a significance level of 10% adjusted *p*-value and log_2_ fold change (log_2_FC) ≥ 1. **B.** Venn diagram illustrates the number of DE genes that are unique to or shared between the *Spi*^+^ and *Tpi*^+^ states relative to uninfected controls (*Ctrl)*. **C.** Heat maps display the DE genes with putative PM-associated and immunity-related functions that are shared between the *Spi*^+^ and *Tpi*^+^ datasets. Fold-change values are represented as a fraction of the average normalized gene expression levels from age-matched *Spi*^+^ or *Tpi*^+^ versus uninfected control flies (*Ctrl*). The heat maps (dendrograms) were clustered using Euclidean distance calculation methods in R-package software. The clusters were manually separated into various categories.

We next evaluated the DE genes from the *Spi^+^* and *Tpi*^+^ groups to understand the functional impact of each microbe infection on host physiology. We began by investigating the potential functions of the 38 genes (Fig. 1B), of which 33 were upregulated and five were downregulated in both *Spi^+^* and *Tpi*^+^ groups compared to control (*Ctrl)* group. Of note, among the induced genes, eight encoded proteins associated with the function of the peritrophic matrix (PM) in tsetse flies. These included four *peritrophins*, *chitin synthase* 2 (*chs*2), a chitin binding protein (*CG34282)*, *chitinase* 2, and *chitin deacetylase* 8 (Fig. 1C and Fig. S2A). Tsetse fly PM is composed of a chitinous matrix embedded with glycoproteins, serving as a barrier that separates the gut lumen from the epithelia. This structure protects the gut cells from harmful compounds present in the bloodmeal as well as from ingested pathogens [46]. The eight induced putative products noted above have previously been described with functions related to PM structure and development in *Gmm* [24] as well as with chitin production or degradation processes [47–49]. Increased expression of genes encoding PM proteins, including Peritrophins (*Pro1* and *Pro3/trypsin),* has also been observed in trypanosome infected *Gpg* [50]. We previously noted compromised PM function(s) in *Gmm* infected with trypanosomes in the midgut and salivary glands as well as in newly eclosed young adults following a bloodmeal supplemented with trypanosomes [51, 52]. Because our results indicated variations in the expression of products involved in PM functions in *Spi^+^* individuals relative to *Ctrl*, we investigated the status of PM integrity by using entomopathogen *S. marcescens* in a fly survival assay as we previously reported [24, 51–53]. Typically, flies with compromised PM survive longer as their gut epithelia can detect the presence of *S. marcescens* and elicit an immediate and robust response that eliminates the pathogen before it causes a fatal systemic infection [24]. We did not observe a statistically significant difference in host survival between *Spi^+^* and *Ctrl* individuals (Fig. S2B), suggesting that the presence of *Spiroplasma* does not influence the structural integrity of PM in the tsetse fly. It is possible that the increased expression of PM associated genes in *Spi*^+^ and *Tpi*^+^ individuals may represent a process of enhanced PM degradation in the presence of pathogens and hence the higher levels of PM associated gene expression to produce more PM proteins to accommodate this process.

In addition to PM-related functions, we detected immunity-related genes that were induced in both *Spi^+^* and *Tpi*^+^ transcriptome datasets compared to *Ctrl*, including C-type lectin, *mucin-5AC* (GFUI18_012666), and *nitric oxide synthase* (NOS) (Fig. 1C, Fig. 2SA). These products have been shown to be part of the immune response to trypanosome infections in *Gmm* [41], with the reactive oxygen species nitric oxide (NO) generated by NOS, exhibiting potent trypanocidal activity [54, 55]. It is possible that the increased expression of these immune molecules in the presence of *Spiroplasma* may contribute to the enhanced parasite refractoriness observed in *Spi*^+^ individuals.

**Fig. 2.**
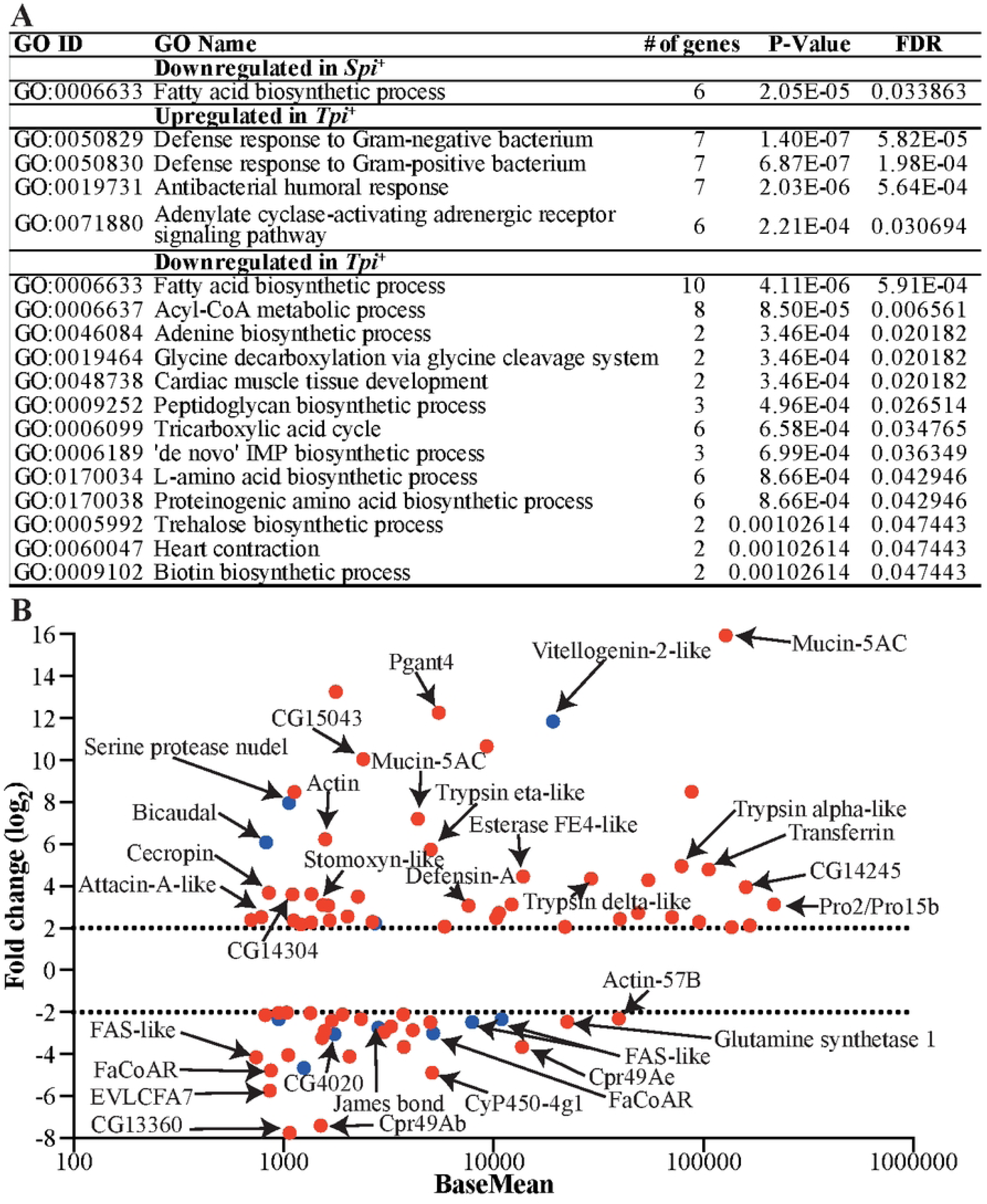
Functional classification of DE-pathways and abundant DE genes unique in *Spi*^+^ or *Tpi*^+^ state. **A.** Gene ontology (GO) enrichment analysis for biological processes was conducted with unique DE genes. **B.** Functional annotations are provided for some of the most abundant and highly DE genes unique to each microbidal infection state, defined by normalized counts ≥ 700 and (log_2_FC) ≥ 2. In this represenetation, red dots indicate genes that are DE in the *Tpi^+^*dataset, while blue dots indicate genes that are DE in the *Spi^+^* dataset.

We also detected DE genes unique to each infection: 31 in the *Spi^+^* group and 951 in the *Tpi*^+^ group (Table S2. Sheet 3 and 4). To gain a comprehensive knowledge on the biological process(es) affected by *Spiroplasma* or trypanosome infections, we subjected the microbe-specific DE genes (log_2_FC ≥ 1) to gene ontology (GO) enrichment analysis using Blast2GO software. The GO enrichment analysis of downregulated putative products in both the *Spi*^+^ and *Tpi*^+^ datasets revealed a shared pathway linked to metabolic processes, particularly lipid biosynthesis,. This was supported by the presence of several genes encoding fatty acid synthases (FAS), fatty acyl CoA reductases, and Acyl-CoA binding protein (Fig. 2A, Fig. S3A). Interestingly, the genes associated with the suppressed fatty acid biosynthesis pathway (GO:0006633) were among the most abundant and highly DE (normalized counts ≥ 700 and log_2_FC ≥ 2) products in both the *Spi*^+^ and *Tpi*^+^ datasets (Fig. 2B; Fig. S3A). A reduction of lipid availability in the gut could negatively impact host fitness and may also reduce pathogen survival. Despite the decreased expression of lipid-associated genes, we found three transcripts including *carnitine O-palmitoyltransferase* I, *fatty acyl-CoA reductase wat like*, and *James bond* that were upregulated in the *Tpi*^+^ dataset (Fig. S3B). Additional biological processes significantly suppressed in the *Tpi*^+^ state included the acyl-CoA metabolic process, the tricarboxylic acid cycle, and amino acid metabolism (Fig. 2A).

### Trypanosomes, but not *Spiroplasma* infections, induce canonical immune pathways in *Gff*

Among the genes induced by trypanosomes were those encoding products associated with immunity and adenylate cyclase signaling pathways (Fig. 2A). In fact, some of the most abundant and highly DE genes (normalized counts ≥ 700, log_2_FC ≥ 2) in the *Tpi*^+^ group included components of the Immune Deficiency (Imd) pathway, mucins, serine proteases, and redox balance associated proteins (Fig. 2B, Fig. S3B). In contrast, these immunity related genes were not induced in the *Spi*^+^ group. The induced immunity-related genes in the *Tpi*^+^ group included two peptidoglycan recognition proteins (PGRPs) -*PGRP-*3 and *PGRP-LA-* along with various antimicrobial peptides (AMPs) such as *Attacin A like*, three *Cecropins*, *Defensin* A and Toll (*Toll-like receptor 7*), defense protein l(2) 34Fc, mucins, C-type lectin 37Db and two phenoloxidase 2. We also identified two genes encoding Homeobox family transcription factors, *Wingless* and *Araucan,* associated with the Wingless signaling pathway (Fig. S3B). In *Gmm,* this pathway has been implicated in the regulation of expression of host *microRNA-275*, which in turn modulates PM-associated gene expression during the trypanosome colonization process [51]. Further gene families upregulated by trypanosomes included trypsins, six serine proteases (SPs), and two serine protease inhibitors (SPIs) linked with insect immunity (Fig. S3C and D). SPs and SPIs play critical roles in modulating the expression of immune pathways (Toll and Imd) by regulating the activation of specific effectors following exposure to an infectious agent [56]. Such regulation ensures that the impact of protease-activated cascades remains localized in time and space [57]. Finally, we detected genes associated with detoxification processes in the *Tpi*^+^ group, including nine *cytochrome P450* (*CYPs*) transcripts, six of which were induced and three reduced. In the *Spi*^+^group, we detected three *CYPs* that were DE, with two significantly induced and one reduced relative to controls (Fig. S3E). The increased expression of *CYPs* in both the *Spi*^+^ and *Tpi^+^* datasets may suggest a protective response to the heightened oxidative stress caused by the presence of trypanosomes and *Spiroplasma*.

### Impact of *Spiroplasma* and/or trypanosome infection on *Gff* lipid metabolism

Our transcriptomic data indicate that infection with *Spiroplasma* and/or trypanosomes negatively impacts the expression of several genes associated with fatty acid biosynthesis, suggesting decreased synthesis and thus low levels of lipids critical for host physiology. We previously demonstrated that pregnant female *Gff* infected with *Spiroplasma* had significantly lower levels of circulating triacyl glyceride (TAG) compared to age-matched, pregnant *Spiroplasma*-negative flies. These flies exhibited reduced fecundity [58], likely as the result of low levels of circulated TAG that make up an important component of tsetse milk [59]. Trypanosomes also scavenge nutrients from their environment, including lipids for metabolism and structural integrity. This nutrient competition may affect critical physiological processes of the host, such as reproductive fitness as previously reported in *Gmm* [60]. With this in mind, we repeated the experiment to investigate the impact of trypanosome infection and, *Spiroplasma* plus trypanosome infections (*Spi*^+^/*Tpi*^+^) on circulating TAG levels in virgin female *Gff*. We compared TAG levels in the hemolymph of two-week-old virgin *Spi*^+^, *Tpi*^+^ and *Spi*^+^/*Tpi*^+^ females with their age-matched *Ctrl* females. Our results showed that virgin *Ctrl* females had higher levels of circulating TAG in their hemolymph (33.75±1.005 μg/μl) compared to their age-matched *Tpi*^+^ (27.72±1.438 μg/μl, *p*=0.0001), *Spi*^+^ (24.90±1.316 μg/μl, *p*<0.0001) and *Spi*^+^/*Tpi*^+^ (19.59±1.438 μg/μl, *p*<0.0001) counterparts (Fig. 3A). In addition, we found that *Spiroplasma*-negative (*Ctrl*) *Gff* males presenr with significantly more (11.56±1.316 μg/μl) circulating TAG than do males infected with the endosymbiont (6.78±1.316 μg/μl, *p*=0.0023) (Fig. 3B). These findings suggest that competition for this critical nutrient impairs host physiology, but this process may also potentially limit pathogen survival and density, as they too rely on host lipids [61, 62].

**Fig. 3.**
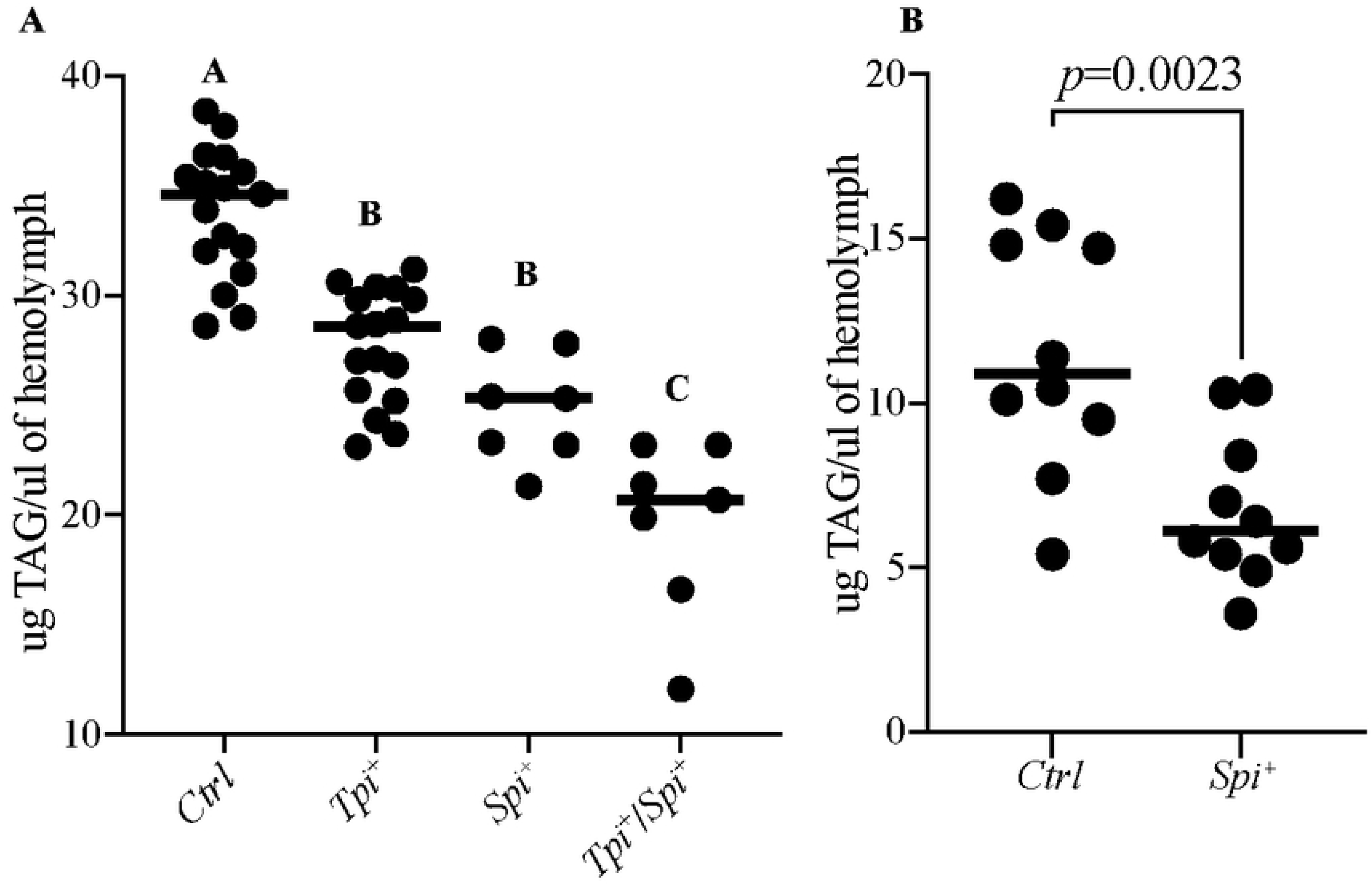
Impact of *Spiroplasma* and/or *Trypanosome* Infection on Host Lipid Level. **A.** Levels of triacylglyceride (TAG) circulating in the hemolymph of virgin female *Gff* were measure for Control (*Ctrl), Spi*^+^, *Tpi*^+^ and *Spi^+^Tpi^+^* groups. **B**. The amount of TAG in the hemolymph of *Spi*^+^ and Control (*Ctrl)* male *Gff* flies. Statistical significance was determined using ANOVA for virgin female flies and unpaired t-test for male flies. Letters A, B and C indicate significant differences (*p*<0.05) in TAG levels, while the the same letter indicates no significant difference (*p*>0.05).

### A newly discovered AMP is expressed in the tsetse fly species in the Palpalis subgenera

One of the most abundant and highly DE genes in the *Tpi^+^* dataset was annotated as *Stomoxyn-like* (GFUI18_001176 or GFUI020894-RA), hereinafter referred to as *GffStomoxyn* (Fig. 2B). The ortholog of *Gff*Stomoxyn was identified in related Diptera, including *S. calcitrans* (stable fly, designated as *Scal*Stomoxyn), *Lucilia sericata* (green blowfly) and *Hermetia illucens* (black soldier fly) [63–65]. The 207 bp *GffStomoxyn* gene encodes a 68 amino acid (aa) pre-pro-mature peptide, which is composed of a 23 aa signal peptide at the N-terminus, followed by pro-mature peptides of 43 aa, similar in organization to the previously described Stomoxyns (Fig. 4A). BLASTP homology searches of the 43 aa *Gff*Stomoxyn mature peptide against proteome databases of *S. calcitrans* and *Musca domestica* (housefly) identified two orthologs in *S. calcitrans,* (SCAU016907; E-value=8.70E−10 and SCAU016937; E-value=0.000632 annotated as *Scal*Stomoxyn 2 and *Scal*Stomoxyn, respectively) and a single ortholog in *M. domestica* (MDOA008330; E-value=1.42E-13 annotated as *Stomoxyn-like* but hereinafter referred to as *Mdom*Stomoxyn). We further searched for the *stomoxyn* locus in other tsetse species using the WGS data from *Gpp* in the Palpalis subgroup, *Gmm* and *Gpd* in the Morsitans subgroup, *Gau* in the Austenina subgroup, and *Gbr* in the Fusca subgroup. This search revealed a single ortholog in *Gpp* (GPPI027903; E-value=1.48E−40), while no orthologs were identified in the other four tsetse species. Additionally, we searched the non-redundant (nr) protein sequence database [66] and relevant literature for putative Stomoxyns. This search identified one ortholog in the flesh fly *Sarcophaga bullata* (DOY81_004902), the previously reported one in *Lucilia sericata* (XP_037825072.1; [64]), and two in the Australian sheep blowfly *L. cuprina (*XP_023308701.2; KAI8119624.1) (Fig. 4A). Interproscan analysis of all identified peptides confirmed their classification within the Stomoxyn protein family. Furthermore, our search also revealed Stomoxyn orthologs in several other Diptera, including *Episyrphus balteatus* (XP_055851874.1) and *Eupeodes corollae* (XP_055904620.1) hoverfly species and the black soldier fly (*H. illucens*) (XP_037911389.1; XP_037913598.1) [65].

**Fig. 4.**
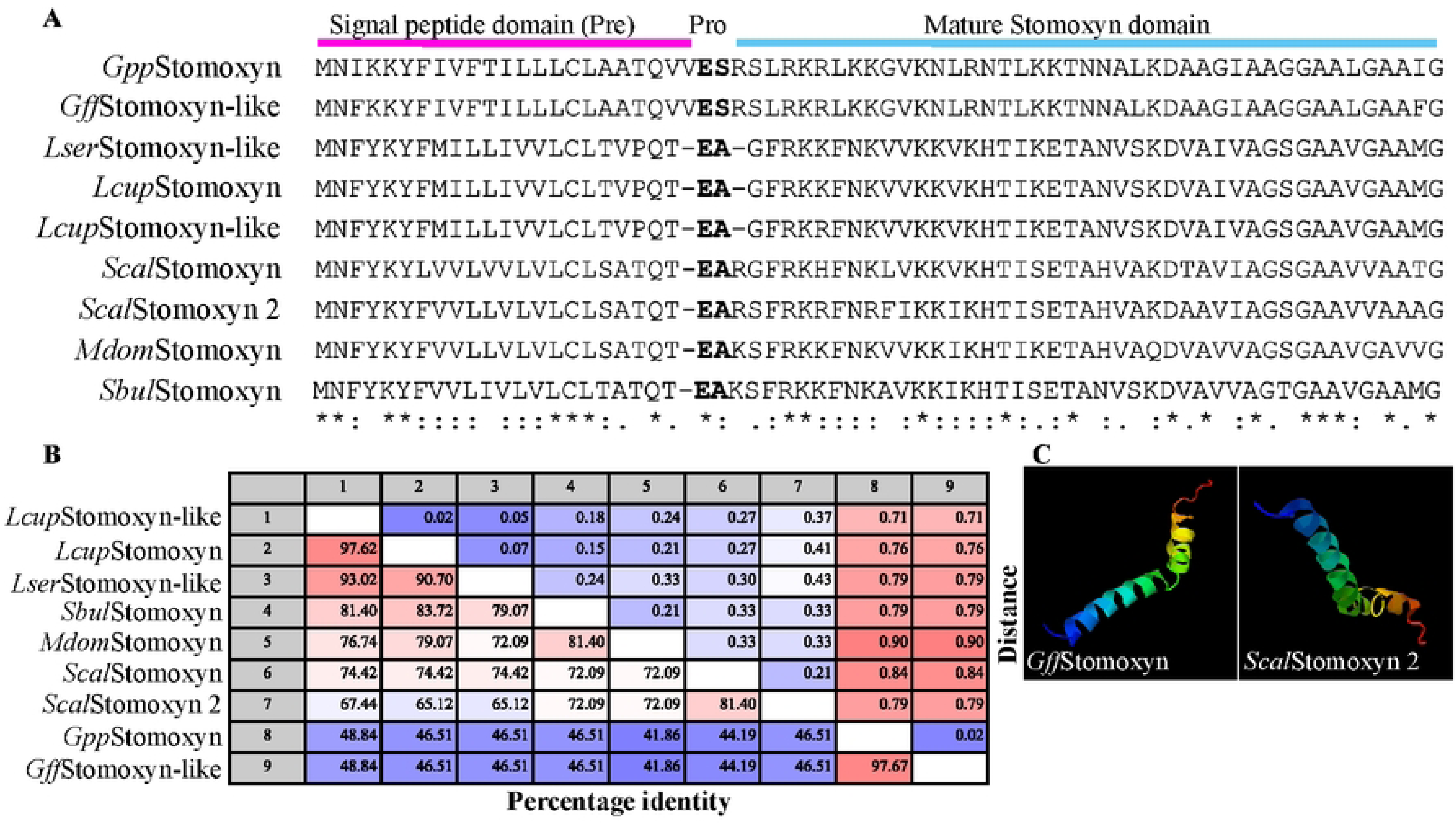
Molecular characterization of *Gff*Stomoxyn. **A.** Sequence alignment of putative Stomoxyn peptides was performed, including sequences from *S. calcitrans* (*Scal*Stomoxyn; SCAU016937 and *Scal*Stomoxyn 2; SCAU016907), *Gff (Gff*Stomoxyn-like; GFUI18_001176), *Gpp (Gpp*Stomoxyn*; GPPI027903*)*, M. domestica* (*Mdom*Stomoxyn-like*; MDOA008330*)*, L. cuprina* (*Lcup*Stomoxyn; KAI8119624.1 and *Lcup*Stomoxyn-like; XP_023308701.2)*, S. bullata* (*Sbul*Stomoxyn; DOY81_004902) and *L. sericata* (*Lser*Stomoxyn-like; XP_037825072.1). The alignment indicates the pre-pro-mature domains of the full predicted peptides, with identical residues indicated by and asterisk (*). Conservative substitutions are marked with a colon (:) and semi-conservative substitutions are marked by a period (.). **B.** Distance matrix analysis was conducted with the mature peptide domains of Stomoxyns from different Diptera. **C.** The tertiary structure of the mature *Gff*Stomoxyn and *Scal*Stomoxyn 2 peptide was predicted by I-TASSER.

Multiple sequence alignment of the putative Stomoxyn peptides revealed a conserved structure comprised of pre-pro-mature domains. Distance matrix analysis of the mature peptide domains showed high identity among the homologs, with *Lcup*Stomoxyn and *Lcup*Stomoxyn-like sharing 97% identify. In contrast, the corresponding domains in the *S. domestica* orthologs, *Scal*Stomoxyn and *Scal*Stomoxyn 2, exhibited only 81% identity. Orthologs within closely related species showed high amino acid identity, such as *Gff* and *Gpp* at 97.67% and *L. sericata* and *L. cuprina* at 93.02% (Fig. 4B). The amino acid identity within the mature peptide domain of Stomoxyns between *M. domestica* and *S. calcitrans* was 72%, while *Gff*Stomoxyn and *Gpp*Stomoxyn exhibited 46% identity with the orthologs from *M. domestica* and *S. calcitrans* (Fig. 4B). Phylogenetic analysis of mature stomoxyns’ amino acid sequences indicate that the *Glossina* stomoxyn-like loci cluster together and are sister to the stomoxys clade, derived from other muscoid flies (Fig. S4A). This stomoxyn phylogeny coincides with host nuclear phylogenies [67], suggesting that further sampling across *Calyptrate* taxa would yield additional stomoxyn-like peptides. I-TASSER prediction of the putative tertiary structure for mature *Gff*Stomoxyn and *Scal*Stomoxyn 2 indicate that they are composed of double α-helices and β-folds (Fig. 4C), similar to the structure reported for *Scal*Stomoxyn [68].

We analyzed the genomic context flanking the *Stomoxyn* loci in *Gff* and *Gpp* and compared their synteny using WGS data available for other *Glossina* species, where no orthologs were identified (Fig. S4B). This analysis revealed several syntenic loci located immediately upstream and downstream of the *Stomoxyn* locus in both *Gff* and *Gpp*. Notably, several loci *GffStomoxyn* were retained on the same genomic scaffolds in *Gpd*, *Gau* and *Gbr*, while such syntenic loci were largely absent from *Gmm* WGS data. Additionally, a BLASTP search of the pre-pro-mature *Gff*Stomoxyn sequence against the putative proteomes of different tsetse species did not yield any significant hits, further suggesting that this protein-coding sequence is absent in those species. To rule out the possibility that the absence of the *Stomoxyn* locus was due to inadequate genome annotation, we performed *de novo* transcript assembly from midgut RNA-seq datasets available for *Gmm* and *Gpd* [51, 69] using Trinity software [70]. We then performed BLASTN searches with the *stomoxyn* sequences collated from *Gff, Gpp*, and other Diptera against our transcriptome-based assembly. This search also did not yield significant hits, further confirming the absence of this locus in *Gmm, Gpd*, *Gau* and *Gbr*.

We next investigated the presence of the *stomoxyn* locus in several *Gff* individuals obtained from distinct populations in North-West Uganda, as well as in *Gpg* from West Africa*, Gbr* and *Gau* from South Africa and *Gau* and *Gpd* from Kenya. We also analyzed this locus in a colony of *Gpp* that was distinct from the one used for the WGS analysis. PCR-based amplification of the *stomoxyn* loci - which included 5’ and 3’ UTRs, two exons and one intron – followed by sequence analysis of the products revealed over 99% identity at the nucleotide level between field collections and flies from laboratory lines. This high level of identity was also observed among *Gff, Gpp* and *Gpg* indicating the close evolutionary relatedness of these fly species (Fig. S4C).

### Analysis of *stomoxyn* expression profile and regulation

We profiled *stomoxyn* expression across various tissues, including cardia, midgut, fat body, female ovary and male testes. Our results indicate that *stomoxyn* is preferentially expressed in the cardia, with significantly lower expression detected in the midgut and other tissues (Fig. 5A). *Stomoxyn* has been similarly reported to be preferentially expressed in the anterior midgut region of *S. calcitrans* [63]. Next, we examined the temporal expression profile of *stomoxyn* and found that midgut transcript levels increased in adults post-eclosion, peaking at 72 h, when newly eclosed flies are mature enough to imbibe their first bloodmeal. *Stomoxyn* levels remained elevated when analyzed 72 h after the first bloodmeal and in the midgut of 15 day-old flies that have consumed multiple bloodmeals (Fig. 5B). We also evaluated the expression of three antimicrobial peptides (AMPs), *stomoxyn*, *cecropin* and *attacin*, in the cardia of 8-day adult flies 72 h after their last bloodmeal. Similarly, we evaluated the expression of these AMPs in the cardia following systemic stimulation by *E. coli* or *per os* stimulation by a BSF *Tbb*-containing bloodmeal. Our results revealed significantly higher levels of *stomoxyn* expression in the cardia compared to the other AMPs and its expression remained high but unresponsive to immune stimuli (Fig. 5C). The levels of both *cecropin* and *attacin* showed a significant increase following *E. coli* challenge, while only *cecropin* was significantly increased following trypanosome challenge (Fig. 5C). We also evaluated the inducible nature of AMP expression in the midgut (gut and cardia) following immune stimulation of teneral *Gff* adults by *per os*challenge with *E. coli*, *S. marcescens,* and BSF *Tbb.* While expression of *cecropin* and *attacin* was significantly induced by *E. coli* and *S. marcescens*, *stomoxyn* expression remained high but unchanged in the midgut (Fig. 5D), similar to our findings in the cardia tissue. We also quantified the *stomoxyn* expression in the midgut of 15-day old adults infected with either *Spiroplasma* (*Spi^+^*) or *Trypanosomes* (*Tpi^+^*) and compared it to uninfected control (*Ctrl)* midguts. Results indicated that *stomoxyn* expression was not influenced by either *Spiroplasma* or trypanosome infection status (Fig. 5E). However, we observed a significant increase in the expression of *attacin* and *cecropin* in *Tpi^+^* relative to the *Ctrl* (Fig. S5A). These results mirror the findings in *Gmm* where it was reported that the expression of both *attacin* and *cecropin* are increased in the trypanosome infected host midguts and have also been shown to have trypanocidal roles [25–27, 71]. Finally, using *Gpg*, another *Palpalis* subgroup tsetse fly, we measured the expression level of *stomoxyn* in the cardia of 8-day old adults and compared it to the AMPs *stomoxyn, attacin* and *cecropin* in non-immune challenged and *Tbb*-immune challenge individuals (Fig. 5F). We found that *stomoxyn* transcript levels were comparable between *Gff* and *Gpg* cardia tissues, and were significantly higher than those of the two AMPs in the unchallenged state. Similarly, we found that neither *stomoxyn* nor *attacin* were induced upon trypanosome challenge in *Gpg* while *cecropin* expression was significantly increased upon trypanosome challenge similar to our findings in *Gff*. Collectively, our results indicate that *stomoxyn* is constitutively expressed at high levels in the cardia of tsetse adults, consistent with the reports of *stomoxyn* in *S. calcitrans* following microbial challenge [63]. Unlike the canonical AMPs, which are inducible in response to pathogen, *stomoxyn* is constitutively expressed in teneral and adult flies.

**Fig. 5.**
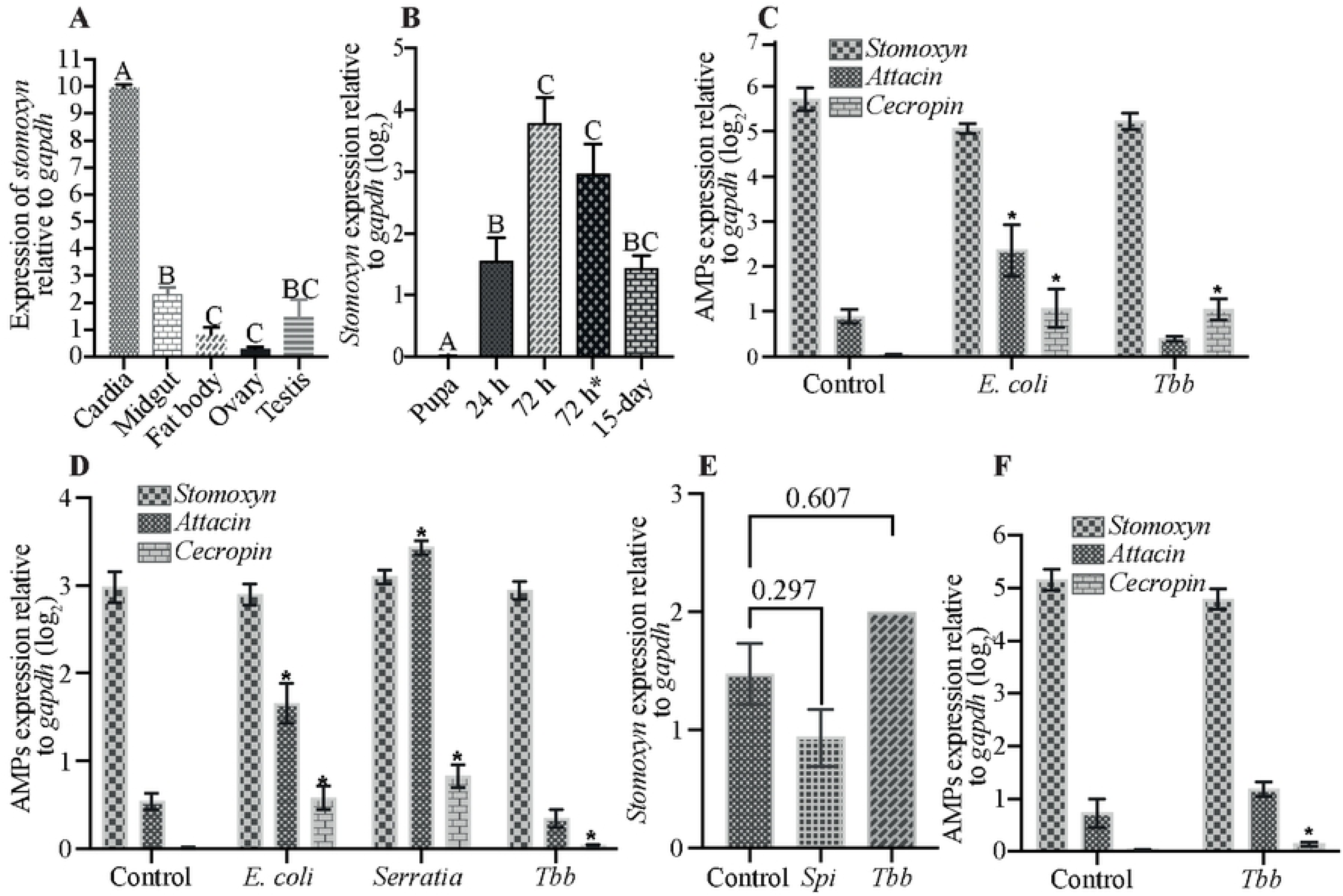
Spatial and Immune Responsive Expression of *GffStomoxyn*. **A.** The qRT-PCR expression profile of *stomoxyn* was assessed from the cardia, midgut, fat body, female ovary, and male testis tissues. Eight biological replicates, each comprising three individual flies, were used in the analysis. Results are presented relative to *gapdh* expression. **B.** Temporal expression of *stomoxyn* was measured from late pupa and whole gut tissue at multiple time points: 24 and 72 hours post-eclosion, 72 hours after first bloodmeal (designated as 72hr*), and in 15 day-old flies analyzed 72 hours after last bloodmeal. Six to ten biological replicates, each comprised of individual midguts, were used in the experiment. Results are presented relative to *gapdh* expression. Letters A, B and C on top of each bar indicate significant differences (*p*<0.05) in *stomoxyn* expression, while the the same letter indicates no significant difference (*p*>0.05). **C.** The expression levels of *stomoxyn, attacin* and *cecropin* in the cardia tissue of 8-day old *Gff* flies 72 hours following microbial challenges with *E. coli* and *Tbb*. This experiment utilized five biological replicates each comprising of three cardia from individual flies. *indicate significantly different gene expression (*p*<0.05) compared to the unchallenged control. **D.** The expression levels of *stomoxyn, attacin* and *cecropin* in the midgut analyzed 24 hours following microbial challenges with *E. coli, Tbb* and *Serratia*. This experiment utilized nine to twelve biological replicates of individual flies. *indicate significantly different gene expression (*p*<0.05) relative to the unchallenged control. **E.** The relative expression levels for *stomoxyn, attacin* and *cepropin* were measured from the in the midguts of trypanosome-infected flies analyzed 15-days post-parasite acquisition and compared to uninfected controls. This experiment utilized eight to twelve biological replicates comprised of individual fly guts. The *p*-values for each comparison are shown above the corresponding bar. **F.** The relative expression levels of *stomoxyn, attacin and cecropin* were determined in the cardia tissue of 8-days old *Gpg* flies 72 hours post-immune challenge with trypanosomes. This experiment utilized five biological replicates each comprising of three cardia from individual flies. *indicate significantly different gene expression (*p*<0.05) relative to the control.

### Antimicrobial activity spectrum of *Gff*Stomoxyn

We next investigated whether *Gff*Stomoxyn exhibits trypanocidal activity similar to that reported for *Scal*Stomoxyn [63]. We had commercially generated synthetic mature *Gff*Stomoxyn, and mature *Scal*Stomoxyn [63] as well as its ortholog, *Scal*Stomoxyn 2. We assessed the microbicidal activity of these three synthetic peptides against *E. coli, Sodalis* (the tsetse fly commensal symbiont) and *Tbb* (both BSF and PCF forms) *in vitro* using minimum inhibition concentration (MIC) assays. The MIC values for *Scal*Stomoxyn, *Scal*Stomoxyn 2 and *Gff*Stomoxyn against *E. coli* were 10 μM, 5 μM and 5 μM, respectively. None of the three peptides exhibited activity against *Sodalis* at concentrations up to the 100 μM (Fig. 6A), consistent with the previously published results for *Scal*Stomoxyn 2 [63, 72]. In contrast, the minimum median inhibitory concentration (MIC_50_) values for *Scal*Stomoxyn, *Scal*Stomoxyn 2 and *Gff*Stomoxyn against BSF trypanosomes were 49.69, 3.51 and 5.06 μM, respectively (Fig. 6B and replicated in Fig. S6A). These results indicate that *Scal*Stomoxyn 2 exhibits significantly stronger killing activity against BSF trypanosomes compared to previously identified *Scal*Stomoxyn (at 3.51 and 49.69 μM, respectively). *Gff*Stomoxyn also demonstrated a high level of trypanolytic activity, with an MIC_50_ of 5.06 μM (Fig. 6B and replicated in Fig. S6A). However, all three peptides were ineffective against PCF trypanosomes at concentrations up to 100 μM (Fig. 6B and replicated in Fig. S6B). Our results indicate that *Gff*Stomoxyn exhibits strong antimicrobial activity against both gram-negative *E. coli* and BSF trypanosomes *in vitro*. Conversely, this peptide is not effective against the endosymbiotic *Sodalis* and insect stage PCF trypanosomes, both of which have coadapted to survive in the insect midgut environement.

**Fig. 6.**
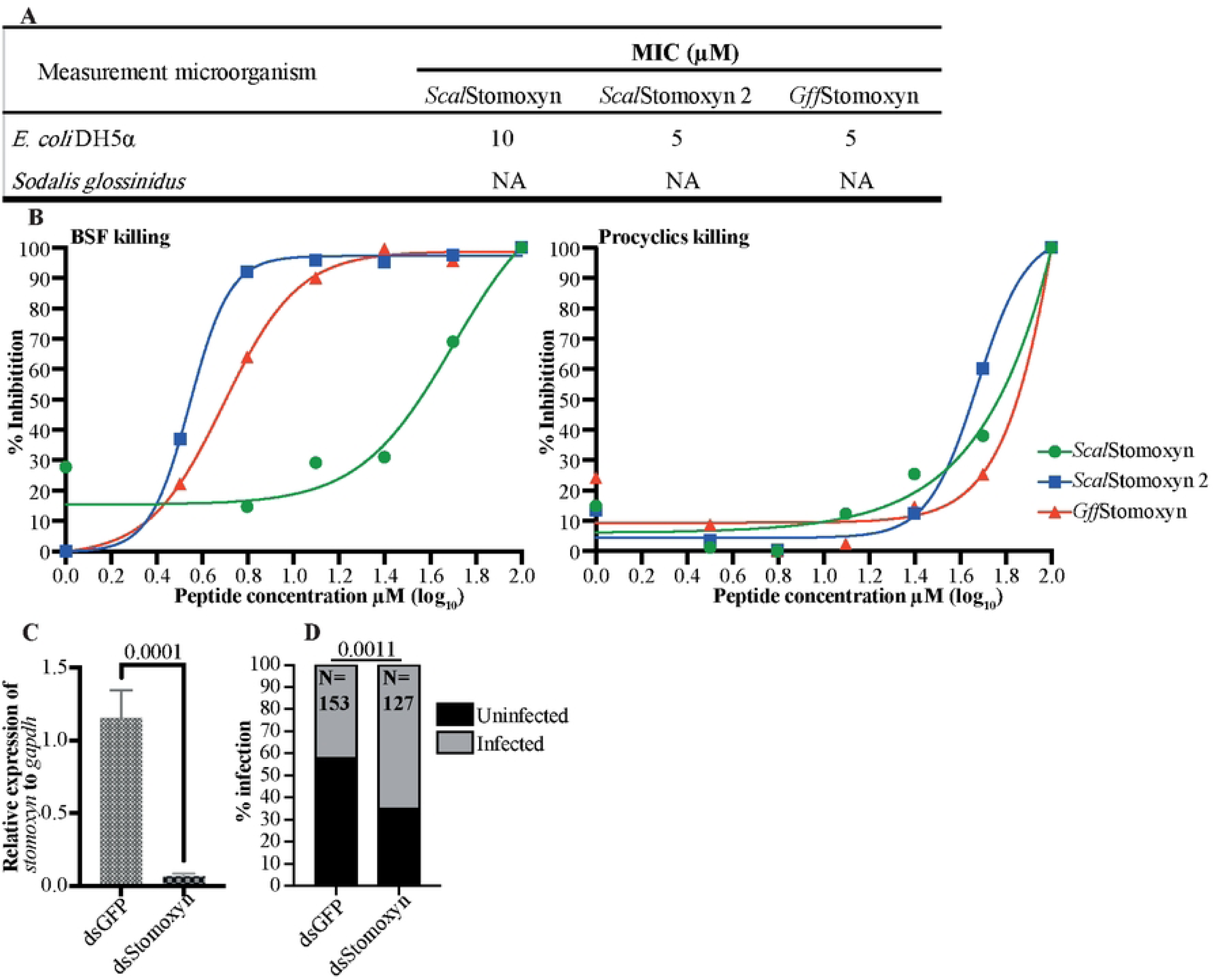
*In vitro* and *in vivo* Bioactivity of Stomoxyn. **A.** *In vitro* antibacterial activity: The antibacterial activity of mature *Scal*Stomoxyn, *Scal*Stomoxyn 2 and *Gff*Stomoxyn peptides was assessed against *E. coli* and *Sodalis* bacteria. Results indicated that *Sodalis* was resistant to Stomoxyn at a maximum concentration of 100μM (NA). **B.** *In vitro* bioactivity against trypanosomes: The bioactivity of mature *Scal*Stomoxyn, *Scal*Stomoxyn 2 and *Gff*Stomoxyn peptide against mammalian bloodstream forms (BSF) and insect stage procyclic form (PCF) trypanosomes. **C.** Gene-silencing validation: The relative expression of *stomoxyn* was measured from dsGFP and dsStomoxyn*-*treated *Gff* flies, showing a significant decrease in expression (*p*<0.0001) in the case of dsStomoxyn*-*treated group. Ten biological replicates, each comprising individual flies, were used to validate silencing efficacy. **D.** Prevalence of trypanosome infections: The prevalence of midgut trypanosome infections in dsRNA-treated *Gff* flies was microscopically analyzed 15 days post-parasite acquisition, with significant difference observed (*p*<0.0011). The total number of flies (N) used in each experimental condition is shown. The experiment included a mixture of male and female flies across two biological replicates, with no significant difference in infection rates between the dsRNA-treated males and females.

### *Stomoxyn* decreases susceptibility of tsetse fly to trypanosome infections

We employed an RNAi based silencing approach to investigate the influence of Stomoxyn peptide on parasite establishment in *Gff*. Double stranded RNAs, dsGFP and dsStomoxyn, were injected into the thoracic haemocoel of teneral adult flies. After 48hs, we confirmed that *GffStomoxyn* expression in dsStomoxyn-treated flies was significantly decreased compared to the dsGFP treated group (*p*<0.0001; Fig. 6C). Forty-eight hours post treatment, both dsGFP and dsStomoxyn flies were offered bloodmeal containing 1×10^6^ BSF trypanosomes and subsequently maintained on normal blood diets. This experiment was replicated twice. On day 15, we analyzed midgut parasite infection prevalence from the two replicates and combined data for analysis. Chi-square analysis revealed that infection rates in the dsStomoxyn group were significantly higher (82/127; 64.56%) compared to the the dsGFP -treated flies (64/153; 41.83%) at df=1, X^2^=10.63, *p* = 0.0011 statistical level of significance (Fig. 6D). These findings indicate that Stomoxyn exhibits trypanocidal activity against BSF trypanosomes *in vivo* in the midgut, similar to the effects observed with synthetic peptides *in vitro*.

## DISCUSSION

Understanding the determinants of vector competence in different tsetse species and subgroups can be complex. These factors may include genetic differences, as revealed by available WGS data, as well as extrinsic factors, such as resident microbiota and the varying ecological niches they occupy. Here, we focused on one of the most important vector species from the genus *Glossina, Gff,* which has a wide distribution in sub-Saharan Africa. Together with closely related species in the Palpalis subgroup, such as *Gpp* and *Gpg, Gff* is responsible for over 90% of Human African Trypanosomiasis (HAT) cases. Although *Gff* are vectors of HAT pathogens as they reside near human settlement and water bodies and are highly anthropophilic, laboratory studies indicate they also exhibit strong resistance to gut colonization and can restrict parasite transmission. Our findings demonstrate one important genetic factor present in *Gff* and other Palpalis subgroup species, the immune peptide Stomoxyn, which has a strong antiparasitic activity *in vitro* against the mammalian BSF trypanosomes acquired through infectious bloodmeal. Functional studies where we reduced the expression of *Gff*Stomoxyn *in vivo* by dsRNA based RNAi treatments resulted in a higher prevalence of parasite infections relative to control groups treated with dsGFP, confirming our *in vitro* results obtained with synthetic *Gff*Stomoxyn peptide. In addition to this genetic factor, our global gene expression profiling of *Spiroplasma-*infected *Gff* individuals suggest that this symbiotic association negatively influences the host metabolic capacity, specifically in lipid biosynthesis. Quantification of the lipid triglycerides in the hemolymph of *Spiroplasma-*infected *Gff* confirmed a reduction, and these lipids which may be important for trypanosome survival as well as for host fecundity. In conclusion, both the *Spiroplasma* endosymbiont mediated metabolic effects that limit critical nutrients, and the trypanocidal Stomoxyn peptide constitutively expressed in the midgut of young adults contribute to the enhanced resistance to parasites observed in these species.

Using a *Gff* line where *Spiroplasma* infections are maintained in approximately 50-60% of adult individuals, we previously demonstrated a negative correlation between the presence of *Spiroplasma* and trypanosome infection success [45]. Similar results were observed in another species within the subgroup Palpalis, *Gt,* collected from Ghana and Burkina Faso [44]. Our comparative transcriptome analyses revealed that infections with *Spiroplasma* symbionts and trypanosome parasites elicit dramatically different host responses in the midgut. While we detected nearly 1000 differentially expressed (DE) genes in response to the trypanosome infection, only about 70 genes were DE in the presence of *Spiroplasma*. This suggests that trypanosomes are recognized as pathogens, inducing a strong immune response, but *Spiroplasma* endosymbiont behaves like a commensal organism, eliciting minimal host response in the midgut. Interestingly, this subdued host response to the presence of *Spiroplasma* in the midgut is contrary to the strong responses we reported in the gonads of tsetse flies where the symbiont affects host reproductive physiology, including sperm viability [58]. However, the enhanced host response in the gonads may reflect the higher density of the endosymbiont present in these tissues relative to the midgut [42]. A common host response to infections with either trypanosome or *Spiroplasma* was a reduction in the expression of genes associated with lipid biosynthesis, suggesting reduced availability of lipids for physiological functions in hosts with microbial infections. Indeed, our TAG measurement assay comparing *Spi*^+^, *Tpi*^+^ and *Spi*^+^/*Tpi*^+^ to the *Ctrl* indicated significantly reduced level of circulating TAG in the hemolymph of virgin microbe-infected female and male *Gff* flies. These findings mirror our previously reported results where we observed lower levels of circulating triglycerides in the hemolymph of *Spiroplasma-*infected pregnant female *Gff* [58]. Such reduction could result from *Spiroplasma* and/or trypanosomes scavenging these metabolites as nutrient sources [61, 73], or from decreased transcriptional activity in lipid biosynthesis pathways. Given that lipids are critical for immunity, reproduction and as a source of energy for both the host and its microbial partners [74, 75], decreased lipid levels can impact tsetse as well as trypanosome physiology and parasite infection success. Similar to what we observed in *Spiroplasma* infected flies, we noted that trypanosome infected *Gff* individuals (*Tpi*^+^), also down-regulate expression of genes associated with fatty acid synthesis and reduced fatty acid biosynthetic processes. In the triatomine bug *Rhodnius prolixus,* infection with *Trypanosoma cruzi* also decreases the expression of several genes associated with lipid biosynthesis [76]. In the *Tpi*^+^ dataset, we detected enhanced expression of *carnitine O-palmitoyltransferase* I (*cpt1*), a gene that encodes for protein that transport fatty acids as acylcarnitines across the outer membrane into the matrix of the mitochondrial for β-oxidation. The increased expression of *cpt1* may represent an adaptive response to produce more acyl-carnitines in order to maintain metabolic homeostasis. In fact, trypanosome forms dwelling in the insect midgut (PCF) have been shown to produce ATP conventionally via β-oxidation in the mitochondria [77, 78], suggesting that they may also utilize the carnitine shuttle for energy production [62, 79, 80]. Collectively our results suggest that both *Spiroplasma* and trypanosome parasites may consume critical nutrients from their host, such as triglycerides, and disrupt metabolic processes essential for host functions, including immunity and reproduction [73, 81].

In the *Tpi*^+^ dataset, the GO analysis of the putatively induced products indicated enrichment of immunity pathways, including *C-type lectins, phenoloxidases, mucins, PGRP, tsetseEP* protein and several AMPs. A similar response to *Tbb* infection was reported in the species *Gmm,* where the expression of many key immune-associated genes was induced in various body compartments [22, 25, 52, 55, 63, 82–85]. The functional roles of some of these immune effectors in trypanosome transmission have been validated through RNAi studies, including IMD, *attacin*, *cecropin* [26, 86], *tsetseEP protein* [31] and *PGRP-LB* [35]. Tsetse species that exhibit greater resistance to trypanosome infections, such as *Gpp* and *Gpd,* express higher levels of Phenoloxidase compared to the susceptible *Gmm* [87]. Studies in *Gmm* indicated that parasite exposure induces *nos* resulting in increased trypanolytic nitric oxide (NO) activity as well as an reactive oxygen intermediate (ROI) and hydrogen peroxide (H_2_O_2_) levels in the cardia organ [54, 55]. These outcomes may mediate molecular communications between local and systemic responses to clear parasite infection. In contrast, *Spiroplasma* infection did not significantly modulate the expression of genes associated with known canonical immune pathways, except for an increase in *nos* levels, which may confer additional parasite resistance to *Spiroplasma*-infected *Spi*^+^ flies. A previous study in *Drosophila melanogaster* also found that *Spiroplasma* infection did not induce epithelial immune responses [88]. This subdued response could result from the absence of a cell wall structure in *Spiroplasma* which contains immune-activating Pathogen-Associated Molecular Patterns (PAMPs) needed to elicit antimicrobial responses [89, 90].

We also report our discovery that *Gff* abundantly expresses the gene encoding Stomoxyn, the orthologue (*Scal*Stomoxyn) first identified in the stable fly, *S. calcitrans.* Stomoxyns are structurally similar to the cecropin family of antimicrobial peptides, which are linear amphipathic in nature with an α-helical structure that lacks cysteine residues. *Gff*Stomoxyn is produced as a pre-pro-mature peptide and has the predicted 3D α-helix structure similar to that of *Scal*Stomoxyn [68]. Our genomic investigations indicate the presence of a single stomoxyn locus in *Gff*, *Gpp, M. domestica* and *L. sericata,* while the genomes of *S. calcitrans, L. cuprina,* and *H. illucens* each contain two copies of this locus, with the homologs showing over 90% identity [64, 65, 91–93]. These homologs may represent recent gene duplication events given that they are more closely related in sequence to one another than they are to their orthologs across taxa, and they appear to be located on the chromosome as tandem copies. Despite multiple investigations, we found that the *stomoxyn* locus is missing from flies in different subgenera of Glossina, including laboratory lines of *Gmm*, *Gpd* and *Gbr* as well as *Gpd, Gau* and *Gbr* natural populations. Prior investigations using cDNA and gDNA of *Gmm* also failed to detect the presence of a *stomoxyn-like* gene in this species [63]. However, the homolog of *GffStomoxyn* is present in the genome of *Gpp* in a region that is sympatric with *Gff*, suggesting a common ancestor. Although WGS data is not available for other species within the Palpalis subgroup, PCR-amplification and sequence analysis indicate that the *stomoxyn* ortholog is also present in *Gpg*. Investigations of *Gff* and *Gpg* obtained from Uganda and Mali have confirmed its presence in natural populations. Hence it appears that *stomoxyn* is a subgroup-specific gene within the genus *Glossina*. Given that the Palpalis subgroup represents a recent expansion in the evolution of *Glossina*, it remains to be determined whether the ancestral lineages lost the s*tomoxyn* locus or if the species within the Palpalis subgroup acquired it post-speciation. Initial analysis of the contigs containing the *stomoxyn* locus for potential flanking mobile element-like sequences did not reveal any candidates that could suggest a potential mechanism of acquisition. Further studies are necessary to investigate how this gene locus was acquired exclusively by this *Glossina* subgroup.

In *S. calcitrans, stomoxyn* is constitutively expressed in the anterior midgut, and due to its trypanocidal activity, has been suggested to confer trypanosome resistance in this hematophogus insect, which is phylogenetically related to *Glossina* [63]. We also found *GffStomoxyn* to be preferentially and constitutively expressed in the cardia organ of *Gff* in the anterior gut. Since the expression of insect AMPs, such as *attacin* and *cecropin*, is known to be induced upon detection of PAMPs by the immune system shortly after infection, we investigated whether *GffStomoxyn* expression might also be immune-responsive. In our transcriptome data, we noted an increase in *GffStomoxyn* in the *Spi*^+^ and *Tpi*^+^ datasets, although the induction by *Spiroplasma* was below the level of significance (*p*=0.1268, adjusted p=0.584 for *Spi*^+^; *p*= 0.0318, adjusted p =0.089 for *Tpi*^+^). Similarly, our qPCR experiments did not validate a significant induction of *GffStomoxyn* expression in the midgut following *per os* challenge with *E. coli* or trypanosomes, nor in *Spiroplasma*-infected *Gff*. This lack of validation could be partly due to the high variability observed in expression levels across the different biological replicates analyzed. Such variation could result from the spatial and temporal nature of *GffStomoxyn* expression, which is preferentially localized to the small cardia organ in the anterior midgut and exhibits increasing expression levels over 72 hours post bloodmeal acquisition.

Although the mature *Gff*Stomoxyn peptide exhibits only 45-50% identity with other Stomoxyns, it has retained its microbicidal and trypanocidal activity. In our *in vitro* assays with synthetic peptides, *Gff*Stomoxyn had a stronger microbicidal activity than the previously described *Scal*Stomoxyn and was similar to that of the *Scal*Stomoxyn2 ortholog discovered in the *S. calcitrans* genome. However, varying purity levels of the synthetic peptides may have contributed to this discrepancy and as such requires further investigations. Functional studies with Stomoxyns from *S. calcitrans, H. illucens* and *L. sericata* also indicated broad-spectrum activity against Gram-negative and Gram-positive bacteria, as well as fungi [63–65, 72, 93]. Our studies with synthetic *Gff*Stomoxyn and *Scal*Stomoxyn2 indicated that both peptides are effective against the mammalian BSF trypanosomes with similar LD_50_ values, but are not effective against the insect stage procylic (PCF) parasites. In addition, *Gff*Stomoxyn showed high antibacterial activity against *E. coli* but not against the *Sodalis* endosymbiont, mirroring earlier reports of *Scal*Stomoxyn activity [72]. Experiments assessing the density levels of the symbionts *Sodalis* and *Wigglesworthia* in *Spiroplasma*-infected *Gff* also showed no reductions compared to control *Gff* [58]. Functional experiments in which we successfully reduced *GffStomoxyn* in the cardia of teneral flies via RNAi found a significant increase in midgut trypanosome infection prevalence in the ds*Gff*Stomoxyn group relative to control dsGFP groups, confirming that *Gff*Stomoxyn is active *in vivo*. We hypothesize that *Gff*Stomoxyn may play a crucial role and be one of the factors that interfere with parasite viability shortly after acquisition in an infected-bloodmeal. This interference likely occurs before the BSF parasites transform into PCF cells within a few days, at which point they are rendered resistant to the trypanolytic actions of this peptide, even at high concentrations.

In conclusion, our results indicate that the reduced vector competence associated with *Gff* and even more so by *Spiroplasma* infections could arise from both unique genomic characteristics and competitive nutritional dynamics between the vector and its symbiont. The reduced lipid production in *Spiroplasma*-infected *Gff* likely limits the availability of essential nutrients required to maintain metabolic hemostasis for both the vector and trypanosome parasites. In addition, host immune responses to *Spiroplasma*, including the increased expression of NOS, can elevate oxidative stress levels in the midgut, resulting in lethal effects on BSF parasites acquired in the infected bloodmeal. Finally, the high levels of trypanolytic Stomoxyn peptide constitutively produced in the cardia organ in the anterior midgut may enhance the clearance of BSF parasites early in the infection process before parasites transform into more resistant PCF cells and colonize the midgut organ. Given that *Sodalis,* the tsetse fly commensal endosymbiont, is resistant to the microbicidal actions of Stomoxyn, it may be feasible to develop genetically modified rec*Sodalis* lines that express Stomoxyn. Ability to colonize tsetse flies with such rec*Sodalis* symbionts through a paratransgenic approach could facilitate the introduction of parasite-resistant flies into field populations, effectively replacing their naturally susceptible counterparts and thereby reduce disease transmission.

## Material and Methods

### Biological material

*Glossina fuscipes fuscipes* (*Gff*) pupae were obtained from the Joint FAO/IAEA IPCL insectary in Seibersdorf, Austria and reared at Yale University insectary at 26°C with 70-80% relative humidity and a 12 hour light:dark photo phase. This *Gff* line carries the heterogenous symbiotic infection, with about 50% of flies being negative (uninfected) for *Spiroplasma*. Newly eclosed teneral *Gff* females were challenged with 2×10^6^ *Trypanosoma brucei brucei* BSF parasites (*Tbb* RUMP 503 strain) in their first bloodmeal and were hereafter maintained on defibrinated bovine blood every 48 hours through an artificial membrane feeding system [94] for 15 days. Another group of the *Gff* flies received normal bloodmeals, were maintained under the same conditions and used for uninfected controls. Whole midguts (MG, comprising cardia and midgut) were dissected 48 hours after the last bloodmeal in Phosphate-buffered-Saline-Glucose (PSG) buffer (pH 8.0) and trypanosome infection status was microscopically determined using a Zeiss Axiostar Plus light microscope (Carl Zeiss Microscopy GmbH, Jana, Germany). All dissected tissues were stored individually at −80°C for RNA extractions. Legs corresponding to individual flies were also collected and kept separately at −80°C for genomic DNA (gDNA) production and determination of *Spiroplasma* infection status.

### Genomic DNA extraction and *Spiroplasma* infection determination

The legs obtained above from individual flies were used to extract gDNA using the Monarch Genomic DNA purification kit following manufacturer’s protocol (New England BioLabs, MA, USA). A *Gff* specific *alpha*-*tubulin* primer set was used for gDNA quality control, and *Spiroplasma*-specific 16s rDNA primer set was used to determine the infection status of the bacterium in each fly. The *Spiroplasma* locus was amplified using touchdown PCR as previously described [45]. Based on the PCR results for *Spiroplasma* infection status, the samples were pooled for downstream analysis as either *Spiroplasma* uninfected (*Ctrl*: control negative for *Spiroplasm*a) or *Spiroplasma* infected (*Spi^+^*; positive only for *Spiroplasma*). Flies that were provided a bloodmeal supplemented with *Tbb* and microscopically found to be infected as described above were similarly screened for the presence of *Spiroplasma*. Of the trypanosome positive flies, only those that were *Spiroplasma* negative were used for downstream analyses (*Tpi*^+^, positive only for trypanosomes). All primer sequences and PCR-amplification conditions used as in Table S1.

### RNA extraction, RNA-seq library preparation and Bioinformatics analysis

Total RNA was extracted from the midgut of all adult individuals using TRIzol and subsequently treated with Turbo-DNase to eliminate contaminating DNA following the protocol described by the manufacturer (Thermo Fisher Scientific Inc., CA, USA). Elimination of DNA from the RNA was confirmed by PCR amplification using *Gff-*specific *alpha*-tubulin and glyceraldehyde-3-phosphate dehydrogenase (*gapdh*) primer sets (Table S1). RNA quantity and quality was determined using an Agilent 2100 Bioanalyzer RNA Nano chip (Agilent, Palo Alto, CA, USA). RNA from two individuals was pooled per biological replicate and four, five and five biological replicates representing *Ctrl*, *Spi^+^*, and *Tpi*^+^ groups, respectively were obtained.

RNA-seq libraries were prepared using the NEBNext Ultra RNA Library Prep Kit for Illumina (New England BioLabs, Inc., MA, USA) according to the manufacturer’s protocol. The individual libraries were barcoded for Illumina HiSeq 2500 sequencing system (Illumina, Inc., CA, USA) and paired-end sequenced (100 bases) at Yale Center for Genome Analysis (YCGA, New Haven, CT). Read files are deposited in the National Center for Biotechnology Information (NCBI) archive, BioProject ID PRJNA1112339.

Raw RNA-seq reads were checked for quality using FASTQC and parsed through Trimmomatic for quality trimming [95]. For analysis, the *Gff* 2018 reference genome version 63 was obtained from VectorBase [96] (https://www.vectorbase.org/). The quality filtered reads were aligned using STAR (v2.3) [97] and ‘htseq-count’ function in the HTSeq (v0.11.2) [98] was used to count the number of reads mapped per gene with the intersection-nonempty mode. Using counts data from HTSeq, correlation between replicates within each condition was evaluated by calculating Pearson’s correlation coefficient (*r)* value. Only genes that had ≥ 10 reads mapping in at least 50% of the biological replicates for each experimental condition were used for downstream analysis. DESeq2 [99] was then used to determine differentially expressed (DE) genes between the control and the infected groups. Genes were considered significantly DE if they exhibited log_2_ fold change either ≥ 1 or ≤ −1 at a *p* value < 0.05 and an adjusted *p value* < 10% [84, 100]. Gene ontology (GO) enrichment analysis of the significantly DE genes was performed using the Blast2GO software [101] using the blastx [102] algorithm at significance threshold of 1×10^−3^ to search against non-redundant (NR) NCBI protein database. An enrichment analysis via Fisher’s exact test at an FDR, p-value ≤ 0.05 in Blast2GO was conducted to determine overexpressed GO terms, relative to the entire *Gff* transcriptome.

### Investigation for Peritrophic Matrix integrity

To assess the effect of *Spiroplasma* infection on the PM integrity, we made use of a host survival assay following infection of the *Gff* line described above with *Serratia marcescens* strain db11 [24, 51]. Briefly, newly eclosed teneral flies were provided a bloodmeal supplemented with *S. marcescens* (1×10^3^ CFU/mL). All flies were then maintained on normal bloodmeals and the number of fly deaths were recorded every 48h. Dead flies at each collection date were kept individually at −80°C for subsequent assessment of *Spiroplasma* infection status via the PCR amplification assay described above. This assay was repeated twice.

### Hemolymph Triacylglycerol (TAG) assay in *Spiroplasma* of Trypanosome infected *Gff*

Hemolymph (3 μl/fly) was collected as described in [103] from two-week-old *Ctrl*, *Spi*^+^, *Tpi*^+^ and *Spi*^+^/*Tpi*^+^ virgin female *Gff*, centrifuged (4°C, 3000xg for 5 minutes) to remove bacterial cells, diluted 1:10 in PBS containing 1.2 μl/ml of 0.2% phenylthiourea (to prevent hemolymph coagulation) and immediately flash frozen in liquid nitrogen [58]. Hemolymph TAG levels were quantified colorimetrically by heating samples to 70°C for 5 min followed by a 10 min centrifugation 16,000xg. Five μl of the supernatant was added to 100 μl of Infinity Triglycerides Reagent (Thermo Scientific) and samples were incubated at 37°C for 10 min. Absorbance was measured at 540nm using a BioTek Synergy HT plate reader as described [58, 104]. Male *Gff* flies *Ctrl* and *Spi*^+^ were also included in the experiment and treated the same way. All *Gff* sample spectra data were compared to that generated from a triolein standard curve (0–50 μg, 10 μg increments). Analysis of Variance (ANOVA) was employed in the analysis of the amount of TAG circulating in the hemolymph of *Ctrl*, *Spi*^+^, *Tpi*^+^ and *Spi*^+/^*Tpi*^+^ female and t-test for circulating TAG in *Gff* male flies’ hemolymph using GraphPad Prism software v.7 (GraphPad Software, La Jolla CA, USA).

### Genomic context and structural analysis of Stomoxyn peptides

To confirm the VectorBaseDB [96] annotation of *Gffstomoxyn-like* gene (GFUI18_001176 or GFUI020894-RA), Blastp [102] homology searches were performed using the putative proteomes for *Stomoxys calcitrans* and *Musca domestica* [92, 105]. This analysis revealed one ortholog in *Glossina palpalis palpalis* (*Gpp*Stomoxyn, GPPI027903), the previously reported Stomoxyn peptide in *S. calcitrans* annotated as *ScalStomoxyn* (SCAU016907) [63] as well as an additional ortholog (annotated as *ScalStomoxyn 2,* SCAU016907) and a single ortholog in *M. domestica* (annotated as *Stomoxyn-like* MDOA008330). Search of published literature and analysis of NR database identified the presence of other orthologs in related Diptera, including *Sarcophaga bullata* (DOY81_004902)*, Lucilia sericata* (XP_037825072.1), two in *Lucilia cuprina (*XP_023308701.2 and KAI8119624.1)*, Episyrphus balteatus* (XP_055851874.1)*, Eupeodes corollae* (XP_055904620.1) and two in *Hermetia illucens* (XP_037911389.1; XP_037913598.1). Full length protein sequence alignment of all significant Blast hits using clustalW [106] and InterProScan analysis [107] were performed to determine conserved Stomoxyn structural and functional domains. RAxML-NG with LG+G8+F model, with 25 parsimony, 25 random starting trees and 1000 bootstraps [108] was used to develop phylogenic tree based on MAFFT [109] sequence alignement of the mature Stomoxyn domain. The resulting tree was visualized with FigTree (http://tree.bio.ed.ac.uk/software/figtree/) and comparatively analyzed with that present in the *Gff* and *Gpp* proteome. The tertiary structure of the Stomoxyn peptides were predicted using the web based I-TASSER program Full length protein sequence alignment of all significant Blast hits using clustalW [106] and InterProScan analysis [107].

In order to understand the evolutionary history of this locus across *Glossina*, we obtained the WGS data available for the different tsetse species in VectorBaseDB, and compared the regions flanking the *Gff* and *Gpp Stomoxyn* loci with the same regions from *Gmm*, *Glossina pallidipes (Gpd)*, *Glossina austeni* (*Gau*) and *Glossina brevipalpis* (*Gbr*), which lack *GffStomoxyn* ortholog. To do this, genes sequences flanking the *Gff*Stomoxyn (GFUI18_001176) in *Gff* supercontig JACGUE010000004 were compared with those in *Gpp* KQ080227 supercontig that contains *GppStomoxyn* via Blastp searches [102]. Similarly, Blastp was used to query the putative proteomes of other *Glossina* species for the presence of *Gff*Stomoxyn orthologs and the supercontigs where the ortholog exist in the genome.

The presence of the *stomoxyn* locus was also investigated from natural fly populations, Genomic DNA was prepared from *Gff* individuals collected in 2018 from the Albert Nile river drainage in Northwest Uganda (Amuru district: Gorodona (GOR; 3°15’57.6“N, 32°12’28.8”E), Okidi (OKS; 3°15’36.0”N, 32°13’26.4”E), and Toloyang (TOL; 3°15’25.2”N, 32°13’08.4”E)), *Glossina palpalis gambienses* (*Gpg*) collected from Bado Souleymane Mali (15°10’0”N, 7°31’0”W), *Gbr* and *Gau* from Hells Gate South Africa, *Gau* and *Gpd* from Shimba Hills Kenya. Genomic DNA was also prepared from a colony of *Gpp* maintained in Burkina Faso that is distinct from the colony in Seirbersdorf Vienna used for the WGS analysis and *Gff* colony from Seirbersdorf Vienna. The gDNA PCR primers (Table S1) were designed to amplify the entire genomic sequence/region of *stomoxyn* in these species. When PCR products were obtained, they were analyzed by Sanger sequence and aligned using CLC Main Workbench (CLC bio, Cambridge, MA) against *Gff* and *Gpp stomoxyn-like* genomic loci from VectorBase.

### Synthesis of Stomoxyn peptides

The predicted mature sequences of three Stomoxyn peptides were commercially synthesized and procured from ABClonal Science (500 West Cummings Park, Woburn, MA). These included the two homologs of Stomoxyn peptide from *S. calcitrans,* the previously described Stomoxyn (*Scal*Stomoxyn) [63] and Stomoxyn 2 (*Scal*Stomoxyn 2) discovered here from the *S. calcitrans* WGS data, and *Gff*Stomoxyn described here. The synthesized peptide sequences were RSLRKRLKKGVKNLRNTLKKTNNALKDAAGIAAGGAALGAAFG (*Gff*Stomoxyn), RSFRKRFNRFIKKIKHTISETAHVAKDAAVIAGSGAAVVAAAG (*Scal*Stomoxyn 2) and RGFRKHFNKLVKKVKHTISETAHVAKDTAVIAGSGAAVVAATG (*Scal*Stomoxyn). Purity of these synthetic peptides were 85.181%, 89.177% and 90.635%, for *Scal*Stomoxyn, *Scal*Stomoxyn 2 and *Gff*Stomoxyn respectively.

### *In vitro* antibacterial and anti-trypanosomal activity of Stomoxyn peptides

The antimicrobial activity of the synthetic Stomoxyn peptides was tested against *E. coli*, tsetse endosymbiont *Sodalis,* and BSF and PCF trypanosomes. For antimicrobial activity, an overnight *E. coli* culture was inoculated into fresh medium and grown to logarithmic phase (OD_600_ 0.3-0.4 at 37°C to yield about 1.5×10^8^ cells/ml). A 1:10 dilution of this culture was used to test the killing activity of different Stomoxyn peptide concentrations. Stock solutions (1 M) of synthetic peptides were prepared in water and 2-fold serial dilutions were made corresponding to 10 μM down to 1.25 μM. The *E. coli* culture was inoculated with synthetic peptides (0, 1.25, 2.5, 5 and 10μM concentrations) for testing the minimal inhibitory concentration (MIC). Cultures were grown for 1 h at 37°C and plated in duplicate on LB plates which were incubated overnight to measure colony forming units. For *Sodalis* killing assays, the bacteria were cultured on brain-heart infusion agar supplemented with 10% bovine blood as previously described [110] and grown in liquid cultures to OD_600_ 0.3-0.4 at 25°C. The synthetic peptides (0μM, 10μM, 20μM, and 100μM concentrations) were prepared and applied as described above for *E. coli.* The experiment was repeated three times with each synthetic peptide and microorganism.

The anti-trypanosomal activity of Stomoxyn was determined using Alamar Blue assay as described [63, 111] with slight modifications. Briefly, 500 BSF *Antat 1.1* 90:13 *T. b. brucei* cells in 50 μl were added to 96-well microtiter plates containing 50 μl of culture medium with a 2-fold serially diluted *Stomoxyn* peptide [112]. Top and low peptide concentration were evaluated at 100 and 3.125 μM respectively. The test for each concentration was performed in triplicates. As controls, culture media was included for baseline fluorescence as well as parasites without the Stomoxyn peptide. After 72 h of incubation at 37°C, 10 μl of Alamar Blue was added to each well, and the plates were incubated for another four hours. The plates were then read using BioTek cytation1 imaging reader (Agilent Technologies Inc., Santa Clara, CA, USA) at an excitation wavelength of 530 nm and an emission wavelength of 590 nm [63]. The acquired data was analyzed to produce sigmoidal inhibition curves and to determine IC_50_ (median drug concentration inhibiting 50% of fluorescence development) values. The Stomoxyn percentage inhibition was calculated using the formula: % inhibition=100[1-(X-MIN)/(MAX-MIN)] where: X was fluorescence of the sample, MIN fluorescence of the control (media without parasites) and MAX fluorescence of the positive control (culture parasites without drug) [113]. Stomoxyn activity for the PCF *Tbb* RUMP 503 was determined following the same protocol above using cells grown in SDM-79 medium at 28°C [114].

### Analysis of *GffStomoxyn* expression

Analysis of spatial expression of *GffStomoxyn* was performed on the cardia, midgut, fat body and ovary tissues from females, and testis tissue from males dissected from 10-day old *Gff*. For temporal expression analysis, whole midgut including cardia was dissected from teneral adults 24 and 72 h post eclosion and from adults 72 h after their first bloodmeal. Fifteen-day old flies that have taken multiple bloodmeals were similarly included in the experiment. To determine the immune responsive profile of *GffStomoxyn* expression, three separate experimental groups of adult teneral flies were provided with bloodmeals supplemented with 1,000 colony forming units (CFU) of *E. coli, S. marcescens* strain db11, and 1×10^6^ BSF *T. b. brucei* (RUMP 503) per ml, respectively. Flies that did not take the bloodmeal were removed from the experiment and the midguts of remaining flies were dissected 24 h later for RNA extraction. Additionally, we exposed adult non-teneral flies (flies that have taken two bloodmeals), *Gff* and *Gpg* to 1000 CFU of E. coli or 1×10^6^ BSF *T. b. brucei* and removed flies that did not take the bloodmeal. The cardia of these flies were dissected 72 h later (n=5 biological replicates each containing three cardia) for RNA extraction. Control groups that received only normal bloodmeals were similarly prepared.

For analysis of trypanosome infected adults, *Gff* were given 1×10^6^ BSF *T. b. brucei* in their first bloodmeal and maintained on normal uninfected bloodmeal for 15 days. Forty-eight hours post last bloodmeal, the midgut of these flies dissected and the presence of trypanosome parasites in the midgut microscopically determined using a Zeiss Axiostar Plus light microscope (Carl Zeiss Microscopy GmbH, Jana, Germany). Trypanosome infected midguts were individually kepts at - 80°C till RNA extraction. Another group of uninfected control *Gff* flies were similiarly treated by being maintained on uninfected bloodmeal. Legs corresponding to individual flies were also collected and kept separately at −80°C for gDNA extraction and determination of *Spiroplasma* infection status of these flies as described above. Only flies determined to be *Spiroplasma* negative (*Ctrl*) and *Spiroplasma* infected (*Spi*^+^) in the control group and those that were only trypanosome positive (*Tpi*^+^) in the exposed group were used for RNA extractions., subsequent cDNA synthesis and further downstream analysis.

Total RNA was prepared (and DNase treated) from these samples as described above. cDNA was synthesized with oligo-dT primers and random hexamers using the iScript cDNA synthesis reaction kit (Bio-Rad, Catalog No. 170–8891) according to the manufacturer’s protocol. Real time quantitative PCR (qRT-PCR) *GffStomoxyn* and other genes (Table S1) were performed in technical duplicate for each sample. The expression level of *GffStomoxyn* was evaluated between tissues and/or experimental conditions by qRT-PCR analysis using *Gff gapdh* as internal control. All qRT-PCR results were thus normalized to tsetse *gapdh*, quantified from each biological replicate in the tissues or experimental condition. The qRT-PCR data was analyzed using Relative Expression Software Tool (REST)-384 version 2 software [115].

### RNAi gene silencing and trypanosome infection prevalence

*Green fluorescent protein* (*gfp*) and *GffStomoxyn* specific dsRNAs were prepared using the MEGAscript High Yield T7 transcription kit (Ambion, Huntingdon, UK) and gene specific dsRNA primers (Table S1). The PCR products dsGFP and ds*Gff*Stomoxyn were sequenced to confirm their specificity for the gene of interest. To test the efficacy of gene silencing, groups of teneral male and female flies 48 h post eclosion were intrathoracically microinjected with 5 µg ds*Gff*Stomoxyn or dsGFP in 2 µl nuclease-free water and the midguts were dissected 48 h post treatment. RNA was extracted from the dissected midguts and analyzed by qRT-PCR amplification for *GffStomoxyn* expression from each treatment group as described above. For trypanosome infection effects, two groups of teneral flies treated with ds*Gff*Stomoxyn or dsGFP as above were provided 1×10^6^/ml BSF *Tbb* in their first bloodmeal administered 48 h post dsRNA treatments. Flies that did not feed were discarded, and all remaining flies were subsequently maintained on normal diets. Fifteen days post-trypanosome challenge, all surviving flies were dissected, and their midguts microscopically examined for the presence of parasite infections. Chi-square was used to compare proportions of trypanosome infections between dsGFP and ds*Gff*Stomoxyn treatment groups. Gene silencing and trypanosome infection experiment was done twice.

## Acknowledgements

This work was generously supported with funding to Serap Aksoy from the Li Foundation, Ambrose Monell Foundation, and NIH/NIAID (RO1AI068932). We are thankful for support from the United Nations, International Atomic Energy Association sponsored Coordinated Research Project entitled ‘Improvement of Colony Management in Insect Mass-Rearing for SIT Applications’. We thank Mr. Sidiya Mbodj for technical support with the tsetse husbandry and Aksoy laboratory members for their critical comments on the manuscript.

## Supplementary data

**Fig. S1. Overview of the *Gff* Transcriptome Study.**

**A.** Table summarizes results obtained from different biological replicates across three conditions, *Ctrl*, *Spi*^+^ and *Tpi^+^*. Condition: Ctrl: *Spiroplasma* and trypanosome negative midgut; *Spi*^+^: *Spiroplasma* positive midgut; *Tpi*^+^: trypanosome positive midgut; BRep: Biological Replicate; UMR: Number of Uniquely Mapped Reads to the *Gff* genome (*Gff*_genome-2018_ver 63); No. Trans: Number of expressed transcripts, defined as those with normalized read coverage ≥ 10 in at least 50% of the biological replicates per condition.

**B.** Heat map showing the Euclidean distances between biological replicates, calculated from the regularized log transformation of the data, provides insights into the similarities and differences among the conditions.

**Fig. S2. Differentially Expressed Shared Transcripts and Peritrophic Matrix Integrity.**

The Heat map denotes the fold changes of differentially expressed (DE) genes that are shared between the *Spi^+^* and *Tpi^+^* states, according to their putative functions. Fold-change values are expressed as a fraction of the average normalized gene expression levels from age-matched *Spi*^+^ or *Tpi*^+^ relative to the control *Ctrl*. The heat maps (dendrograms) were generated using Euclidean distance calculation combined with ward.D clustering methods within the R-package software. The clusters were manually separated into two categories: PM and Immunity Functions.

**B.** Effect of *Spiroplasma* infection on Peritrophic Matrix integrity. The survival of flies was monitored every 48 hours following a *per os* treatment of teneral adult flies with *Serratia marcescens*, administered 72 hours post-eclosion. At time of death, the *Spiroplasma* infection status of each fly was evaluated using our diagnostic assay. The Kaplan-Meyer survival curves illustrate the fly survival over time for *Spiroplasma*-uninfected flies (blue) and *Spiroplasma*- infected flies (red). This experiment was conducted twice, with no significant differences observed between the two experiment (Data not shown).

**Fig. S3. Heatmap representation of unique differentially expressed (DE) transcripts.**

The heatmaps depict the fold changes of unique DE transcripts across various functional categories, comparing infected transcriptomes and the uninfected control. *Spi*+: *Spiroplasma* infected, *Tpi*+: trypanosome infected; FC: fold change indicate the degress of change in expression levels relative to uninfected controls; 1: differentially expressed transcripts; NDE: transcripts that are not differentially expressed.

**Fig. S4. Genomic Characterization of Stomoxyn locus.**

**A.** Phylogenetic tree of mature Stomoxyn sequences from nine different Diptera species based the Maximum likelihood (ML) model. Sequences used in this analysis were obtained from VectorBase for *S. calcitrans* (*Scal*Stomoxyn; SCAU016937 and *Scal*Stomoxyn 2; SCAU016907), *Gff (Gff*Stomoxyn-like; GFUI18_001176), *Gpp (Gpp*Stomoxyn*; GPPI027903*) and *M. domestica* (*Mdom*Stomoxyn-like*; MDOA008330*), and from NCBI database for *L. cuprina* (*Lcup*Stomoxyn; KAI8119624.1 and *Lcup*Stomoxyn-like; XP_023308701.2)*, S. bullata* (*Sbul*Stomoxyn; DOY81_004902), *L. sericata* (*Lser*stomoxyn-like; XP_037825072.1), *Episyrphus balteatus* (*Ebal*Stomoxyn-like; XP_055851874.1) and *Eupeodes corollae* (*Ecor*Stomoxyn-like; XP_055904620.1). The analysis involved 11 amino acid sequences and 1000 bootstrap replications. **B.** Genomic content surrounding the Stomoxyn-like gene (GFUI18_001176) focusing on the supercontig JACGUE010000004 of the Gff genome assembly (version 63, Vectorbase). The supercontigs available from the other *Glossina* WGS data were also compared. Genes exhibiting synteny among different tsetse species are indicated by arrows with the same color genes that do not shown synteny are presented in gray. The black arrow marks the expected location of the *Stomoxyn* gene in *G. pallidipes, G. austeni* and *G. brevipalpis*. Gene size and spacings are not drawn to scale. Based on the current assembly of the region, only one *Stomoxyn* gene is present in *Gff* and *Gpp,* while it is absent in other *Glossina* species. **C.** Multiple sequence alignment of the Stomoxyn locus from *Gff*, *Gpp* and *Gpg.* The alignment includes genomic PCR product sequences of the *Stomoxyn* locus from flies obtained from laboratory and field populations, confirming the presence and conservation of this locus in the species from Palpalis subgroup. The primers for the PCR amplification were designed to span the entire coding region of the Stomoxyn pre-pro-mature peptide.

**Fig. S5. Expression of *attacin* and *cecropin* in *Tpi^+^* flies 15 days post emergence.**

Flies were exposed to a bloodmeal spiked with BSF *Tbb* in their first meal and fifteen days later, their midguts were dissected 48 hours after their last bloodmeal. A control group of flies were treated similarly but received normal blood meals without any parasites. *Trypanosome* and *Spiroplasma* infection status of each fly was determined as described in materials and methods section. A total of eleven biological replicates were analyzed for *Ctrl* and eight for *Tpi*^+^ groups. The expression of *attacin* and *cecropin* were quantified using RT-qPCR relative to tsetse *gapdh*. Expression levels showed significant difference (*p*=0.001) between the *Tpi*^+^ and *Ctrl* groups.

**Fig. S6. Bioactivity of Stomoxyn Against Trypanosomes. A.** *In vitro* killing activity of rec*Gff*Stomoxyn against mammalian BSF trypanosomes **B.** *In vitro* killing activity of rec*Gff*Stomoxyn against insect specific procyclic PCF trypanosomes. Data shown represent findings from the second replicate experiment showing the potency of *Gff*Stomoxyn against BSF and PCF forms.

**Table S1. PCR primers used in this study.**

**Table S2. Detailed results and analysis of each transcriptome.**

## Reference

1. WHO. Trypanosomiasis, Human African (Sleeping Sickness). 2019 [cited 2023 November 5, 2019]; Available from: www.who.int/news-room/fact-sheets/detail/ trypanosomiasis-human-african- (sleeping-sickness).

2. Holt, H.R., et al., Assessment of animal African trypanosomiasis (AAT) vulnerability in cattle-owning communities of sub-Saharan Africa. Parasit Vectors, 2016. 9: p. 53.

3. Maudlin, I., P.H. Holmes, and M.A. Miles, *The Trypanosomiasis. CAB International*. 2004, Wallingford, UK.: CABI Publishing.

4. Silva Pereira, S., A.P. Jackson, and L.M. Figueiredo, Evolution of the variant surface glycoprotein family in African trypanosomes. Trends Parasitol, 2022. 38(1): p. 23–36.

5. Holmes, P., Tsetse-transmitted trypanosomes--their biology, disease impact and control. J Invertebr Pathol, 2013. 112 Suppl: p. S11–4.

6. Aksoy, S., et al., Human African trypanosomiasis control: Achievements and challenges. PLoS Negl Trop Dis, 2017. 11(4): p. e0005454.

7. Ndeffo-Mbah, M.L., et al., The impact of vector migration on the effectiveness of strategies to control gambiense human African trypanosomiasis. PLoS Negl Trop Dis, 2019. 13(12): p. e0007903.

8. Sutherland, C.S., et al., Seeing beyond 2020: an economic evaluation of contemporary and emerging strategies for elimination of Trypanosoma brucei gambiense. Lancet Glob Health, 2017. 5(1): p. e69–e79.

9. Msangi, A.R., C.J. Whitaker, and M.J. Lehane, Factors influencing the prevalence of trypanosome infection of Glossina pallidipes on the Ruvu flood plain of Eastern Tanzania. Acta Trop, 1998. 70(2): p. 143–55.

10. Matetovici, I., L. De Vooght, and J. Van Den Abbeele, Innate immunity in the tsetse fly (Glossina), vector of African trypanosomes. Dev Comp Immunol, 2019. 98: p. 181–188.

11. Geiger, A., F. Ponton, and G. Simo, Adult blood-feeding tsetse flies, trypanosomes, microbiota and the fluctuating environment in sub-Saharan Africa. Isme j, 2015. 9(7): p. 1496–507.

12. Welburn, S.C., D.H. Molyneux, and I. Maudlin, Beyond Tsetse--Implications for Research and Control of Human African Trypanosomiasis Epidemics. Trends Parasitol, 2016. 32(3): p. 230–241.

13. Moloo, S.K. and S.B. Kutuza, Comparative study on the infection rates of different laboratory strains of Glossina species by Trypanosoma congolense. Med Vet Entomol, 1988. 2(3): p. 253–7.

14. Kennedy, P.G., The continuing problem of human African trypanosomiasis (sleeping sickness). Ann Neurol, 2008. 64(2): p. 116–26.

15. Welburn, S.C., I. Maudlin, and P.P. Simarro, Controlling sleeping sickness - a review. Parasitology, 2009. 136(14): p. 1943–9.

16. Holdsworth, P., et al., World Association for the Advancement of Veterinary Parasitology (WAAVP) second edition: Guideline for evaluating the efficacy of parasiticides against ectoparasites of ruminants. Vet Parasitol, 2022. 302: p. 109613.

17. Cecchi, G., et al., Land cover and tsetse fly distributions in sub-Saharan Africa. Med Vet Entomol, 2008. 22(4): p. 364–73.

18. Rayaisse, J.B., et al., Towards an optimal design of target for tsetse control: comparisons of novel targets for the control of Palpalis group tsetse in West Africa. PLoS Negl Trop Dis, 2011. 5(9): p. e1332.

19. Rayaisse, J.B., et al., Prospects for the development of odour baits to control the tsetse flies Glossina tachinoides and G. palpalis s.l. PLoS Negl Trop Dis, 2010. 4(3): p. e632.

20. Attardo, G.M., et al., Comparative genomic analysis of six Glossina genomes, vectors of African trypanosomes. Genome Biol, 2019. 20(1): p. 187.

21. Miesen, P., J. Joosten, and R.P. van Rij, PIWIs Go Viral: Arbovirus-Derived piRNAs in Vector Mosquitoes. PLoS Pathog, 2016. 12(12): p. e1006017.

22. Hao, Z. and S. Aksoy, Proventriculus-specific cDNAs characterized from the tsetse, Glossina morsitans morsitans. Insect Biochem Mol Biol, 2002. 32(12): p. 1663–71.

23. Rose, C., et al., An investigation into the protein composition of the teneral Glossina morsitans morsitans peritrophic matrix. PLoS Negl Trop Dis, 2014. 8(4): p. e2691.

24. Weiss, B.L., et al., The peritrophic matrix mediates differential infection outcomes in the tsetse fly gut following challenge with commensal, pathogenic, and parasitic microbes. J Immunol, 2014. 193(2): p. 773–82.

25. Hao, Z., et al., Tsetse immune responses and trypanosome transmission: implications for the development of tsetse-based strategies to reduce trypanosomiasis. Proc Natl Acad Sci U S A, 2001. 98(22): p. 12648–53.

26. Hu, C. and S. Aksoy, Innate immune responses regulate trypanosome parasite infection of the tsetse fly Glossina morsitans morsitans. Mol Microbiol, 2006. 60(5): p. 1194–204.

27. Hu, Y. and S. Aksoy, An antimicrobial peptide with trypanocidal activity characterized from Glossina morsitans morsitans. Insect Biochem Mol Biol, 2005. 35(2): p. 105–15.

28. Wang, J., et al., Characterization of the antimicrobial peptide attacin loci from Glossina morsitans. Insect Mol Biol, 2008. 17(3): p. 293–302.

29. Macleod, E.T., et al., Factors affecting trypanosome maturation in tsetse flies. PLoS One, 2007. 2(2): p. e239.

30. MacLeod, E.T., et al., Antioxidants promote establishment of trypanosome infections in tsetse. Parasitology, 2007. 134(Pt 6): p. 827–31.

31. Haines, L.R., et al., Tsetse EP protein protects the fly midgut from trypanosome establishment. PLoS Pathog, 2010. 6(3): p. e1000793.

32. Stiles, J.K., et al., Identification of trypanolysin and trypanoagglutinin in Glossina palpalis sspp*. (*Diptera: Glossinidae). Parasitology, 1990. 101: p. 369–376.

33. Nyambega, B., et al., Lysis of Trypanosoma brucei brucei by Tsetse Trypanolysin. Kenya Journal of Science (B series), 2011. 14: p. 26–34.

34. Osir, E.O., M. Abakar, and L. Abubakar. The role of trypanolysin in the development of trypanosomes in tsetse. in Proceedings of the 25th Meeting of the International Council for Trypanosomiasis Research Control (ISCTRC). 1999. Mombasa, Kenya.

35. Wang, J., et al., Interactions between mutualist Wigglesworthia and tsetse peptidoglycan recognition protein (PGRP-LB) influence trypanosome transmission. Proceedings of the National Academy of Sciences, 2009. 106(29): p. 12133–12138.

36. Abubakar, L., E.O. Osir, and M.O. Imbuga, Properties of a blood-meal-induced midgut lectin from the tsetse fly Glossina morsitans. Parasitol Res, 1995. 81(4): p. 271–5.

37. Abubakar, L.U., et al., Molecular characterization of a tsetse fly midgut proteolytic lectin that mediates differentiation of African trypanosomes. Insect Biochem Mol Biol, 2006. 36(4): p. 344–52.

38. Osir, E.O., L. Abubakar, and M.O. Imbuga, Purification and characterization of a midgut lectin-trypsin complex from the tsetse fly Glossina Iongipennis. Parasitol Res, 1995. 81: p. 276–281.

39. Imbuga, M.O., E.O. Osir, and V.L. Labongo, Inhibitory effect of Trypanosoma brucei brucei on Glossina morsitans midgut trypsin in vitro. Parasitol Res, 1992. 78(4): p. 273–6.

40. Imbuga, M.O., et al., Studies on tsetse midgut factors that induce differentiation of bloodstream Trypanosoma brucei brucei in vitro. Parasitol Res, 1992. 78: p. 10–15.

41. Welburn, S.C. and I. Maudlin, Tsetse-trypanosome interactions: rites of passage. Parasitol Today, 1999. 15(10): p. 399–403.

42. Doudoumis, V., et al., Challenging the Wigglesworthia, Sodalis, Wolbachia symbiosis dogma in tsetse flies: Spiroplasma is present in both laboratory and natural populations. Sci Rep, 2017. 7(1): p. 4699.

43. Mfopit, Y.M., et al., Molecular detection of Sodalis glossinidius, Spiroplasma species and Wolbachia endosymbionts in wild population of tsetse flies collected in Cameroon, Chad and Nigeria. BMC Microbiol, 2023. 23(1): p. 260.

44. Dera, K.M., et al., Prevalence of Spiroplasma and interaction with wild Glossina tachinoides microbiota. Parasite, 2023. 30: p. 62.

45. Schneider, D.I., et al., Spatio-temporal distribution of Spiroplasma infections in the tsetse fly (Glossina fuscipes fuscipes) in northern Uganda. PLoS Negl Trop Dis, 2019. 13(8): p. e0007340.

46. Aksoy, S., Tsetse peritrophic matrix influences for trypanosome transmission. J Insect Physiol, 2019. 118: p. 103919.

47. Yang, X.B., et al., Identification and RNAi-Based Functional Analysis of Four Chitin Deacetylase Genes in Sogatella furcifera (Hemiptera: Delphacidae). J Insect Sci, 2021. 21(4).

48. Khajuria, C., et al., A gut-specific chitinase gene essential for regulation of chitin content of peritrophic matrix and growth of Ostrinia nubilalis larvae. Insect Biochem Mol Biol, 2010. 40(8): p. 621–9.

49. Merzendorfer, H. and L. Zimoch, Chitin metabolism in insects: structure, function and regulation of chitin synthases and chitinases. J Exp Biol, 2003. 206(Pt 24): p. 4393–412.

50. Hamidou Soumana, I., et al., Comparative Genomics of Glossina palpalis gambiensis and G. morsitans morsitans to Reveal Gene Orthologs Involved in Infection by Trypanosoma brucei gambiense. Front Microbiol, 2017. 8: p. 540.

51. Aksoy, E., et al., Mammalian African trypanosome VSG coat enhances tsetse’s vector competence. Proc Natl Acad Sci U S A, 2016. 113(25): p. 6961–6.

52. Vigneron, A., et al., A fine-tuned vector-parasite dialogue in tsetse’s cardia determines peritrophic matrix integrity and trypanosome transmission success. PLoS Pathog, 2018. 14(4): p. e1006972.

53. Yang, L., et al., Paratransgenic manipulation of a tsetse microRNA alters the physiological homeostasis of the fly’s midgut environment. PLoS Pathog, 2021. 17(6): p. e1009475.

54. Bogdan, C., Nitric oxide and the immune response. Nat Immunol, 2001. 2(10): p. 907–16.

55. Hao, Z., I. Kasumba, and S. Aksoy, Proventriculus (cardia) plays a crucial role in immunity in tsetse fly (Diptera: Glossinidiae). Insect Biochem Mol Biol, 2003. 33(11): p. 1155–64.

56. Christophides, G.K., et al., Immunity-related genes and gene families in Anopheles gambiae. Science, 2002. 298(5591): p. 159–65.

57. Bouton, M.C., et al., The under-appreciated world of the serpin family of serine proteinase inhibitors. EMBO Mol Med, 2023. 15(6): p. e17144.

58. Son, J.H., et al., Infection with endosymbiotic Spiroplasma disrupts tsetse (Glossina fuscipes fuscipes) metabolic and reproductive homeostasis. PLoS Pathog, 2021. 17(9): p. e1009539.

59. Benoit, J.B., et al., Adenotrophic viviparity in tsetse flies: potential for population control and as an insect model for lactation. Annu Rev Entomol, 2015. 60: p. 351–71.

60. Hu, C., et al., Infections with Immunogenic Trypanosomes Reduce Tsetse Reproductive Fitness: Potential Impact of Different Parasite Strains on Vector Population Structure. PLoS Neglected Tropical Diseases, 2008. 2(3): p. e192.

61. Herren, J.K., et al., Insect endosymbiont proliferation is limited by lipid availability. Elife, 2014. 3: p. e02964.

62. Poudyal, N.R. and K.S. Paul, Fatty acid uptake in Trypanosoma brucei: Host resources and possible mechanisms. Front Cell Infect Microbiol, 2022. 12: p. 949409.

63. Boulanger, N., et al., Epithelial innate immunity. A novel antimicrobial peptide with antiparasitic activity in the blood-sucking insect Stomoxys calcitrans. J Biol Chem, 2002. 277(51): p. 49921–6.

64. Pöppel, A.K., et al., Antimicrobial peptides expressed in medicinal maggots of the blow fly Lucilia sericata show combinatorial activity against bacteria. Antimicrob Agents Chemother, 2015. 59(5): p. 2508–14.

65. Elhag, O., et al., Screening, Expression, Purification and Functional Characterization of Novel Antimicrobial Peptide Genes from Hermetia illucens (L.). PLoS One, 2017. 12(1): p. e0169582.

66. Clark, K., et al., GenBank. Nucleic Acids Res, 2016. 44(D1): p. D67–72.

67. Bayless, K.M., et al., Beyond Drosophila: resolving the rapid radiation of schizophoran flies with phylotranscriptomics. BMC Biol, 2021. 19(1): p. 23.

68. Landon, C., et al., Solution structures of stomoxyn and spinigerin, two insect antimicrobial peptides with an alpha-helical conformation. Biopolymers, 2006. 81(2): p. 92–103.

69. Bateta, R., et al., Tsetse fly (Glossina pallidipes) midgut responses to Trypanosoma brucei challenge. Parasit Vectors, 2017. 10(1): p. 614.

70. Grabherr, M.G., et al., Full-length transcriptome assembly from RNA-Seq data without a reference genome. Nat Biotechnol, 2011. 29(7): p. 644–52.

71. Boulanger, N., et al., Immunopeptides in the defense reactions of Glossina morsitans to bacterial and Trypanosoma brucei brucei infections. Insect Biochem Mol Biol, 2002. 32(4): p. 369–75.

72. Haines, L.R., R.E. Hancock, and T.W. Pearson, Cationic antimicrobial peptide killing of African trypanosomes and Sodalis glossinidius, a bacterial symbiont of the insect vector of sleeping sickness. Vector Borne Zoonotic Dis, 2003. 3(4): p. 175–86.

73. Parreira de Aquino, G., et al., Lipid and fatty acid metabolism in trypanosomatids. Microb Cell, 2021. 8(11): p. 262–275.

74. Kaczmarek, A., et al., Insect Lipids: Structure, Classification, and Function. Adv Exp Med Biol, 2024.

75. Attardo, G.M. and I.A. Hansen, Editorial: Insights into lipid biology and function in insect systems. Front Insect Sci, 2022. 2: p. 1119577.

76. Sousa, G., S.S. de Carvalho, and G.C. Atella, Trypanosoma cruzi Affects Rhodnius prolixus Lipid Metabolism During Acute Infection. Frontiers in Tropical Diseases, 2021. 2.

77. Zíková, A., et al., The F(0)F(1)-ATP synthase complex contains novel subunits and is essential for procyclic Trypanosoma brucei. PLoS Pathog, 2009. 5(5): p. e1000436.

78. Bochud-Allemann, N. and A. Schneider, Mitochondrial substrate level phosphorylation is essential for growth of procyclic Trypanosoma brucei. J Biol Chem, 2002. 277(36): p. 32849–54.

79. Klein, R.A., J.M. Angus, and A.E. Waterhouse, Carnitine in Trypanosoma brucei brucei. Mol Biochem Parasitol, 1982. 6(2): p. 93–110.

80. Steketee, P.C., et al., Transcriptional differentiation of Trypanosoma brucei during in vitro acquisition of resistance to acoziborole. PLoS Negl Trop Dis, 2021. 15(11): p. e0009939.

81. Smith, T.K. and P. Bütikofer, Lipid metabolism in Trypanosoma brucei. Mol Biochem Parasitol, 2010. 172(2): p. 66–79.

82. Telleria, E.L., et al., Insights into the trypanosome-host interactions revealed through transcriptomic analysis of parasitized tsetse fly salivary glands. PLoS Negl Trop Dis, 2014. 8(4): p. e2649.

83. Lehane, M.J., et al., Adult midgut expressed sequence tags from the tsetse fly Glossina morsitans morsitans and expression analysis of putative immune response genes. Genome Biol, 2003. 4(10): p. R63.

84. Matetovici, I., G. Caljon, and J. Van Den Abbeele, Tsetse fly tolerance to T. brucei infection: transcriptome analysis of trypanosome-associated changes in the tsetse fly salivary gland. BMC Genomics, 2016. 17(1): p. 971.

85. Attardo, G.M., et al., Analysis of fat body transcriptome from the adult tsetse fly, Glossina morsitans morsitans. Insect Mol Biol, 2006. 15(4): p. 411–24.

86. Nayduch, D. and S. Aksoy, Refractoriness in tsetse flies (Diptera: Glossinidae) may be a matter of timing. J Med Entomol, 2007. 44(4): p. 660–5.

87. Nigam, Y., et al., Detection of phenoloxidase activity in the hemolymph of tsetse flies, refractory and susceptible to infection with Trypanosoma brucei rhodesiense. J Invertebr Pathol, 1997. 69(3): p. 279–81.

88. Hurst, G.D., et al., Hidden from the host: Spiroplasma bacteria infecting Drosophila do not cause an immune response, but are suppressed by ectopic immune activation. Insect Mol Biol, 2003. 12(1): p. 93–7.

89. Akira, S., S. Uematsu, and O. Takeuchi, Pathogen recognition and innate immunity. Cell, 2006. 124(4): p. 783–801.

90. Kleinnijenhuis, J., et al., Innate immune recognition of Mycobacterium tuberculosis. Clin Dev Immunol, 2011. 2011: p. 405310.

91. Anstead, C.A., et al., Lucilia cuprina genome unlocks parasitic fly biology to underpin future interventions. Nat Commun, 2015. 6: p. 7344.

92. Olafson, P.U., et al., The genome of the stable fly, Stomoxys calcitrans, reveals potential mechanisms underlying reproduction, host interactions, and novel targets for pest control. BMC Biol, 2021. 19(1): p. 41.

93. Hirsch, R., et al., Profiling antimicrobial peptides from the medical maggot Lucilia sericata as potential antibiotics for MDR Gram-negative bacteria. J Antimicrob Chemother, 2019. 74(1): p. 96–107.

94. Moloo, S.K., An artificial feeding technique for Glossina. Parasitology, 1971. 63(3): p. 507–12.

95. Bolger, A.M., M. Lohse, and B. Usadel, Trimmomatic: a flexible trimmer for Illumina sequence data. Bioinformatics, 2014. 30(15): p. 2114–20.

96. Giraldo-Calderon, G.I., et al., VectorBase: an updated bioinformatics resource for invertebrate vectors and other organisms related with human diseases. Nucleic Acids Res, 2015. 43(Database issue): p. D707–13.

97. Dobin, A., et al., STAR: ultrafast universal RNA-seq aligner. Bioinformatics, 2013. 29(1): p. 15–21.

98. Anders, S., P.T. Pyl, and W. Huber, HTSeq--a Python framework to work with high-throughput sequencing data. Bioinformatics, 2015. 31(2): p. 166–9.

99. Love, M.I., W. Huber, and S. Anders, Moderated estimation of fold change and dispersion for RNA-seq data with DESeq2. Genome Biol, 2014. 15(12): p. 550.

100. Benjamini, Y. and Y. Hochberg, Controlling the False Discovery Rate: A Practical and Powerful Approach to Multiple Testing. J R Stat Soc Ser B, 1995. 57.

101. Gotz, S., et al., High-throughput functional annotation and data mining with the Blast2GO suite. Nucleic Acids Res, 2008. 36(10): p. 3420–35.

102. Altschul, S.F., et al., Basic local alignment search tool. J Mol Biol, 1990. 215(3): p. 403–10.

103. Benoit, J.B., et al., Symbiont-induced odorant binding proteins mediate insect host hematopoiesis. Elife, 2017. 6.

104. Agbu, P., et al., MicroRNA miR-7 Regulates Secretion of Insulin-Like Peptides. Endocrinology, 2020. 161(2).

105. Scott, J.G., et al., Genome of the house fly, Musca domestica L., a global vector of diseases with adaptations to a septic environment. Genome Biol, 2014. 15(10): p. 466.

106. Thompson, J.D., D.G. Higgins, and T.J. Gibson, CLUSTAL W: improving the sensitivity of progressive multiple sequence alignment through sequence weighting, position-specific gap penalties and weight matrix choice. Nucleic Acids Res, 1994. 22(22): p. 4673–80.

107. Jones, P., et al., InterProScan 5: genome-scale protein function classification. Bioinformatics, 2014. 30(9): p. 1236–40.

108. Kozlov, A.M., et al., RAxML-NG: a fast, scalable and user-friendly tool for maximum likelihood phylogenetic inference. Bioinformatics, 2019. 35(21): p. 4453–4455.

109. Katoh, K., MAFFT: a novel method for rapid multiple sequence alignment based on fast Fourier transform. Nucleic Acids Res, 2002. 30.

110. Smith, C.L., et al., Characterization of the achromobactin iron acquisition operon in Sodalis glossinidius. Appl Environ Microbiol, 2013. 79(9): p. 2872–81.

111. Räz, B., et al., The Alamar Blue assay to determine drug sensitivity of African trypanosomes (T.b. rhodesiense and T.b. gambiense) in vitro. Acta Trop, 1997. 68(2): p. 139–47.

112. Baltz, T., et al., Cultivation in a semi-defined medium of animal infective forms of Trypanosoma brucei, T. equiperdum, T. evansi, T. rhodesiense and T. gambiense. Embo j, 1985. 4(5): p. 1273–7.

113. Shapiro, A.B., et al., A high-throughput fluorescence resonance energy transfer-based assay for DNA ligase. J Biomol Screen, 2011. 16(5): p. 486–93.

114. Brun, R. and Schönenberger, Cultivation and in vitro cloning or procyclic culture forms of Trypanosoma brucei in a semi-defined medium. Short communication. Acta Trop, 1979. 36(3): p. 289–92.

115. Pfaffl, M.W., G.W. Horgan, and L. Dempfle, Relative expression software tool (REST) for group-wise comparison and statistical analysis of relative expression results in real-time PCR. Nucleic Acids Res, 2002. 30(9): p. e36.

